# TDP-43 stabilizes transcripts encoding stress granule protein G3BP1: potential relevance to ALS/FTD

**DOI:** 10.1101/2020.09.15.298455

**Authors:** Hadjara Sidibé, Yousra Khalfallah, Shangxi Xiao, Nicolás B. Gómez, Elizabeth M.H. Tank, Geneviève Di Tomasso, Eric Bareke, Anaïs Aulas, Paul M. McKeever, Ze’ev Melamed, Laurie Destroimaisons, Jade-Emmanuelle Deshaies, Lorne Zinman, J. Alex Parker, Pascale Legault, Martine Tétreault, Sami J. Barmada, Janice Robertson, Christine Vande Velde

**Affiliations:** Departments of Neurosciences, Université de Montréal, Montreal, QC, Canada; Departments of Biochemistry, Université de Montréal, Montreal, QC, Canada; CHUM Research Center, Montréal, QC, Canada; Center for Neurodegenerative Disease, University of Toronto, Toronto, ON, Canada; Department of Neurology, University of Michigan, Ann Arbor, MI, USA; Neuroscience Graduate Program, University of Michigan, Ann Arbor, MI, USA; Cellular and Molecular Biology Program, University of Michigan, Ann Arbor, MI, USA; Univeristy of California, San Diego/Ludwig Institute for Cancer Research, San Diego, CA, USA; Sunnybrook Health Sciences Centre

**Author notes:** **Corresponding author & Lead author:** Christine Vande Velde, Ph.D., Department of Neurosciences, Universite de Montreal, CRCHUM-Tour Viger, 900, rue Saint-Denis, R09.442, Montreal, QC, CANADA H2X 0A9, poste 28832.

## Abstract

TDP-43 nuclear depletion and concurrent cytoplasmic accumulation in vulnerable neurons is a hallmark feature of progressive neurodegenerative proteinopathies such as amyotrophic lateral sclerosis (ALS) and frontotemporal dementia (FTD). Cellular stress signalling and stress granule dynamics are now recognized to play a role in ALS/FTD pathogenesis. Defective stress granule assembly is associated with increased cellular vulnerability and death. G3BP1 (Ras-GAP SH3-domain-binding protein 1) is a critical stress granule assembly factor. Here, we define that TDP-43 stabilizes *G3BP1* transcripts via direct binding of a highly conserved *cis* regulatory element within the 3’UTR. Moreover, we show *in vitro* and *in vivo* that nuclear TDP-43 depletion is sufficient to reduce G3BP1 protein levels. Finally, we establish that *G3BP1* transcripts are reduced in ALS/FTD patient neurons bearing TDP-43 cytoplasmic inclusions/nuclear depletion. Thus, our data suggest that, in ALS/FTD, there is a compromised stress granule response in disease-affected neurons due to impaired *G3BP1* mRNA stability caused by TDP-43 nuclear depletion. These data implicate TDP-43 and G3BP1 loss of function as contributors to disease.

## INTRODUCTION

TDP-43 is a ubiquitously expressed, mainly nuclear, RNA/DNA binding protein belonging to the heterogeneous nuclear ribonucleoprotein family (hnRNPs). It is a major regulator of RNA metabolism and is involved in several steps of RNA regulation (mRNA transport, nucleocytoplasmic export, stability, transcription, translation, splicing, etc.) (Taylor *et al*., 2016). Like other hnRNPs, TDP-43 has an N-terminal domain containing a nuclear localization sequence (NLS), an RNA binding domain (RBD) composed of two RNA recognition motifs (RRM1 and RRM2) and a glycine rich C-terminal domain, also known as an intrinsically disordered domain (Taylor *et al*., 2016). While the NLS dictates the localization of TDP-43 in the nucleus, the RRMs are essential for specific interaction with target RNAs, and the N and C-terminal domains serve as platforms to mediate TDP-43 self-assembly and interaction with other protein partners, respectively (Kuo *et al*., 2014; Mompean *et al*., 2016; Afroz *et al*., 2017).

TDP-43 proteinopathies refer to a group of neurological disorders characterized by the pathological accumulation of TDP-43 in the cytoplasm of affected neurons and, less frequently, oligodendroglia (Neumann *et al*., 2006). The fatal neurodegenerative diseases ALS and FTD are included in this group. ALS shares clinical, neuropathological and genetic overlaps with frontotemporal dementia (FTD), the second most common cause of dementia in those <65 years, which is caused by the degeneration of the frontal and temporal lobes (FTLD). In addition to the neuromuscular component of the disease, ~50% of ALS patients exhibit cognitive impairment caused by FTLD, with 15% fulfilling the clinical diagnostic criteria of FTD (Forman *et al*., 2007). TDP-43 nuclear depletion and concomitant accumulation in the cytoplasm, often as a component of skein-like or round inclusions, in motor and cortical neurons is observed in 97% of all ALS cases and 50% of FTD patients (Taylor *et al*., 2016). Given these shared clinical and pathological features, ALS and FTD with TDP-43 pathology are now considered to be part of the same disease spectrum (Taylor *et al*., 2016). However, it remains debated as to whether nuclear depletion of TDP-43 with simultaneous cytoplasmic accumulation induces a loss or gain of function effect, or some combination thereof.

The interplay between genetics and the environment is suspected to play a major role in the development and/or progression of ALS and FTD (Al-Chalabi and Hardiman, 2013). Recent work places stress granules, a core aspect of the integrated stress response that facilitates cellular recovery from adverse environmental exposures, as a critical component of disease pathogenesis (Dewey *et al*., 2010; Boeynaems *et al*., 2017; Mackenzie *et al*., 2017; Sidibé and Vande Velde, 2019). These micron-sized cytoplasmic foci form concomitant with translational arrest following stress exposure and are proposed to protect and/or sort non-translating mRNAs (Anderson and Kedersha, 2006, 2008). The key protein for stress granule assembly is G3BP1 (Ras-GAP SH3-domain-binding protein 1) (Tourriere *et al*., 2003; Humoud *et al*., 2016). We have previously demonstrated that G3BP1 protein and transcript levels are decreased as TDP-43 levels are lowered, which compromises cellular viability post-stress (McDonald *et al*., 2011; Aulas *et al*., 2012; Aulas *et al*., 2015; Khalfallah *et al*., 2018). This TDP-43-induced G3BP1 depletion disrupts stress granule dynamics, which can be rescued by the reintroduction of G3BP1 (Aulas *et al*., 2012). Recent work reinforces that G3BP1 is central to stress granule formation as it is a tunable switch regulating the liquid-liquid phase separation (LLPS) required for stress granule formation (McDonald *et al*., 2011; Aulas *et al*., 2012; Guillén-Boixet *et al*., 2020; Sanders *et al*., 2020; Yang *et al*., 2020). Thus, we reason that defining the molecular mechanism underlying TDP-43 modulation of G3BP1 is essential to understanding the contribution of stress granule biology to ALS and FTD pathogenesis.

Herein, we demonstrate that nuclear TDP-43 depletion, as observed in the neurons of ALS/FTD patients, as well as expression of pathological TDP-43 variants associated with ALS/FTD, impact *G3BP1* mRNA metabolism. Additionally, we have uncovered that while there are two *G3BP1* transcripts encoding the same protein, only one of them is regulated by TDP-43. TDP-43 binds and stabilizes the short *G3BP1* transcript via an evolutionary conserved sequence. Finally, we demonstrate that this transcript is the predominant transcript expressed in adult human motor neurons and *G3BP1* mRNA is decreased in ALS/FTD patient neurons featuring TDP-43 nuclear depletion/cytoplasmic localization. Taken together, our work suggests that compromised stress granule assembly due to the loss of TDP-43 nuclear function in *G3BP1* stability may be relevant to disease pathogenesis.

## MATERIALS AND METHODS

### Constructs

Flag-TDP-43^ΔNLS^siRes was generated using the QuickChange^®^ II Site-Directed mutagenesis kit (Agilent Technologies) on pCS2-Flag-TDP-43^WT^ (Kabashi *et al*., 2010) with the forward primer 5’-CAACTATCCAAAAGATAACGCAGCAGCAATGGATGAGACAGATGC-3’ and its complementary reverse to mutate the NLS, as previously published (Winton *et al*., 2008). Modifications to render TDP-43 plasmids siRNA-resistant were performed with the forward primer 5’-CTTCCTAATTCTAAGCAGAGGCAGGACGAGCCTTTGAGAAGC-3’ and its complementary reverse. The His-G3BP1cDNA-3’UTR construct was subcloned into pTRE-tight which was provided by Dr. Yves Berthiaume (Institut de recherche cliniques de Montreal). His-G3BP1 was generated by PCR with the forward primer coding for His-tag: forward 5’-CTAGGATCCGCAATGCACCACCACCACCACCACGTGATGGAGAAGCCTAGTC CCCTG-3’; reverse 5’-ATCGCAGCTAGCGCACCTGCCGTGGCGCAAGC-3’) and inserted using BamHI and NheI restriction sites. G3BP1 3’UTR was generated using PCR amplification with the following primers: forward 5’-ATCGCAGCTAGCGCGTGAATCTTCATGGATCTTCATGCAG-3’ and reverse 5’-GCACTAATCGATGCAAACAAAACCTTCACCCATCTCAC-3’ and inserted using NheI and ClaI restriction sites. Deletion of nt319-372 (GRCh38/hg38 chr5: 151,804,204-151,804,257 was made using the following phosphorylated primers: forward 5’-5/phos/CTTAAGCAGTTTATAACAGACTGGGGTCATA-3’ and reverse 5’-5/phos/CCTGACCTTTAGTCTTTCACTTCCAATTTTG-3’.

### Cell culture and transfection

HeLa, HepG2 and HEK293 cells were cultured in Dulbecco’s high glucose modified Eagle medium (DMEM, GE Healthcare) supplemented with 10% fetal bovine serum (FBS, Wisent) and 2 mM L-glutamine (Sigma). HeLa Tet-off cells were cultured in Dulbecco’s high glucose modified Eagle medium (DMEM, GE Healthcare) supplemented with 10% tetracycline-free FBS (Wisent) and 2 mM L-glutamine. SH-SY5Y and SK-N-SH cells were cultured in Dulbecco’s high glucose modified Eagle medium/Nutrient Mixture F-12 Ham (DMEM-F12, ThermoFisher Scientific) supplemented with 10% FBS (Gibco), 2 mM L-glutamine and 1% MEM non-essential amino acids (ThermoFisher Scientific). Cells were collected 72 h after transfection with 125 pmol Stealth siRNA using Lipofectamine^®^ 2000 (Invitrogen) and 24h after plasmid transfection using Lipofectamine^®^ LTX and Plus reagent (Invitrogen). siRNA sequences used are: Control (siCTL): #12935-200 (Invitrogen), and TDP-43 (siTDP-43): 5’ AAGCAAAGCCAAGAUGAGCCUUUGA-3’, as previously published (McDonald *et al*., 2011; Aulas *et al*., 2012; Aulas *et al*., 2015; Deshaies *et al*., 2018).

### Human neuron differentiation

Human induced pluripotent stem cells (iPSCs) from a healthy 54 year old female were reprogrammed from fibroblasts via transfection with episomal vectors encoding seven reprogramming factors, as previously described (Yu *et al*., 2011; Tank *et al*., 2018). A polycistronic construct containing a doxycycline-inducible cassette driving expression of the transcription factors neurogenins 1 and 2 (Busskamp *et al*., 2014; Lam *et al*., 2017a), Dendra2-tagged TDP-43 (WT or M337V) and eIF3-iRFP was integrated into the *CLYBL* safe harbour locus on chromosome 13. For creation of TDP-43(M337V), the homologous recombination vector used to insert Dendra2 was modified to include the corresponding mutation in the *TARDBP* ORF (c. 1009A>G). All insertions and base pair mutations were verified by PCR and Sanger sequencing. As described in prior work (Fernandopulle *et al*., 2018; Weskamp *et al*., 2019), integration of this cassette enables rapid, robust and consistent differentiations, and produces homogenous cultures of forebrain-like, excitatory, glutamatergic neurons optimal for subsequent biochemical or genetic studies (Cerbini *et al*., 2015). Cells were harvested in Trizol at DIV10. Lines are verified mycoplasma-free on a monthly basis.

### Immunoblotting

Proteins were extracted with RIPA buffer (150 mM NaCl, 50 mM Tris pH 7.4, 1% Triton X-100, 0.1% SDS, 1% sodium deoxycholate) with protease inhibitors (10 μg/ml leupeptin, 10 μg/ml pepstatin A, 10 μg/ml chymostatin), separated by SDS-PAGE, transferred to nitrocellulose and blocked with 5% powdered milk in PBS-T (137mM NaCl, 2.7mM KCl, 8mM Na2HPO4, 1.5mM KH2PO4, 1% Tween-20). Membranes were incubated with rabbit anti-TDP-43 (10789-2-AP; Proteintech), mouse anti-Actin (69100; MP Biomedicals), and mouse anti-G3BP1 (sc-81940; Santa Cruz) followed by HRP-conjugated secondary antibodies (Jackson Immunoresearch) and labeled with ECL chemoluminescence (ThermoFisher Scientific). Films were quantified by densitometry using Photoshop (Adobe, CS4).

### Immunofluorescence

Coverslips were fixed in 1% formaldehyde/PBS, washed with PBS, permeabilized in 0.1% Triton X-100/PBS, and blocked in 0.1% bovine serum albumin/PBS. Coverslips were incubated with primary antibodies: mouse anti-Flag (F1804; Sigma) and rabbit anti-TDP-43 (10789-2-AP; Proteintech) diluted in 0.1% BSA/PBS; washed once with 0.1% Triton X-100/PBS and then twice with 0.1% BSA/PBS. Coverslips were then incubated with secondary antibodies: donkey anti-mouse Alexa 488 (Jackson Immunoresearch) and donkey anti-rabbit Alexa 594 (Jackson Immunoresearch) diluted in 0.1% BSA/PBS, washed, labelled with TO-PRO-3 iodide (ThermoFisher Scientific), and mounted using ProLong Antifade (ThermoFisher Scientific). Images were collected on a Leica TCS SP5 confocal microscope equipped with 40x (1.25 N.A.) oil objective and the Leica Application Suite imaging software.

### Cell lines RT—PCR and qRT-PCR

RNA was extracted using RNAeasy Minikit (Qiagen) and treated with DNAse I (Qiagen) according to the manufacturer’s instructions. Equal amounts of RNA were reverse transcribed using the QuantiTect Reverse Transcription kit (Qiagen). cDNA was amplified with the following primers: *G3BP1* total forward: 5’-AGCCATACAAACCCTGGTTCC-3’; *G3BP1* total reverse: 5’-ACAACAGTTTTGCTTGGTTGT-3’; *G3BP1* long forward: 5’-CACTGAGGGTCCTCCCGATA-3’; *G3BP1* long reverse: 5’-CCAGACCAATGGAAGCACCT-3’; *GAPDH* forward: 5’-GACAGTCAGCCGCATCTTCT-3’; *GAPDH* reverse: 5’-GCGCCCAATACGACCAAATC-3’;. The QuantStudio 7 Flex Real-Time PCR System (Life Technologies) was used for qPCR. PrimeTime Standard qPCR assays (IDT) were: *GAPDH* (targeting exon 1-2) Hs.PT.39a.22214836; *G3BP1* total (targeting exon 6 and 7) Hs.PT.58.20396264; *G3BP1* long forward: 5’-TCTTACTGGACACTCAACCTTG-3’; *G3BP1* long reverse: 5’-TGCCATAACTTTTGTGACTTCATG-3’, *18S* forward: 5’-CCAGTAAGTGCGGTCATAAG-3’; *18S* reverse: 5’-GGCCTCACTAAACATCCAA-3’. Data was analyzed using the 2^-ΔΔ^Ct method. The genes of interest were standardized to the geometric mean of 2 housekeeping genes (*GAPDH* and *18S*)

### RNA immunoprecipitation

HeLa cells were lysed in 10 mM Tris, pH 7.4, 50 mM NaCl, 0.5% Triton X-100, with protease inhibitors, triturated subsequently through 18G and 25G needle syringes, incubated for 5 min at room temperature, centrifuged at 13 *000g* and the supernatant collected. Aliquots of 10 mg of pre-cleared lysate were immunoprecipitated at 4°C overnight with mouse anti-TDP-43 (H00023435-M01; Abnova) or mouse anti-Flag (F1804; Sigma) pre-bound to Protein G Dynabeads (ThermoFisher Scientific). Immunoprecipitates were treated with DNase (Qiagen) and RNA was recovered with TRIzol (Invitrogen). Equal amounts of RNA were reverse transcribed using the QuantiTect Reverse Transcription kit (Qiagen). cDNA was amplified by standard PCR with the following primers: *G3BP1* forward: 5’-AAGCCCCCTTCCCACTCCAA-3’; *G3BP1* reverse: 5’-TAATCGCCTTCGGGGACCTG-3’ (targeting exon 12); *CAMKII*forward: 5’-CCACAGGGGCTTTAGGAGA-3’; *CAMKII* reverse: 5’-GCTGCTGCCGCTTTTGTA-3’; *HSPA1A* forward: 5’-TGAAGGAGACAGCCGAAAG-3’; *HSPA1A* reverse: 5’-TGCATCAGTATAGAAACAAGGAA-3’.

### Luciferase assays

HeLa cells were transfected with siRNA for 48 h, and then subsequently transfected using FuGENE (Promega) for 24 h with reporter plasmids including the promotor and 3’ untranslated region (UTR) of human *G3BP1* (SwitchGear Genomics) and relevant controls, as recommended by the manufacturer (for promoter assay: *GAPDH* and a vector containing random sequence, R01; for 3’ UTR: *GAPDH, HDAC6* and vector containing random sequence, R03). The LightSwitch Assay (Promega) reagents were added according to the manufacturer’s instructions and luciferase activity was assessed with a Synergy H4 Hybrid Multi-Mode Microplate reader (Biotek).

### RNA stability assay

HeLa Tet-off cells were subjected to siRNA using Lipofectamine 2000 (Invitrogen) for 48 h then transfected for 24 h with 1 ug His-G3BP1 3’UTR constructs using Lipofectamine LTX and Plus reagent (Invitrogen). To determine mRNA stability, HeLa cells were treated with 1 μg/ml doxycycline (Sigma) for 2, 4, and 6 h prior to RNA extraction with the RNeasy Minikit (Qiagen). Equal amounts were reverse transcribed via the QuantiTect Reverse Transcription kit (Qiagen), according to the manufacturer’s instructions. The QuantStudio 7 Flex Real-Time PCR System (Life Technologies) was used for qPCR. PrimeTime Standard qPCR assays (IDT) for His-G3BP1 forward: 5’-ACGTGATGGAGAAGCCTAGT-3’; reverse: 5’-TGACATCACTTTCCTGTGGATTT-3’, *18S* forward: 5’-CCAGTAAGTGCGGGTCATAAAG-3’; *18S* reverse: 5’-GGCCTCACTAAACCATCCAA-3’.

### C. elegans RT–PCR

Some strains were provided by the CGC, which is funded by NIH Office of Research Infrastructure Programs (P40 OD010440). RNA from N2 and *tdp-1*(*ok803*) strains was extracted using TRIzol (Invitrogen) and chloroform (Fisher Scientific). Equal amounts of RNA were reverse transcribed using the QuantiTect Reverse Transcription kit (Qiagen). cDNA was amplified with the following primers: *tdp-1* forward 5’-AAAGTGGGATCGAGTGACGAC-3’; *tdp-1* reverse 5’-GACAGCGTAACGAATGCAAAGC-3’; *gtbp-1* forward 5’-TGCCTCGCAAAATGGCAATC-3’; *gtbp-1* reverse 5’-AGGAAAAAGGAGAACGGAAAGC-3’; *actin-3* forward 5’-GTTGCCGCTCTTGTTGTAGAC-3’; *actin-3* reverse 5’-GGAGAGGACAGCTTGGATGG-3’. Data was analyzed by densitometry.

### RNA–pull down

Lysates were prepared in pulldown buffer (10 mM Tris pH 7.4, 50 mM NaCl, 0.5% Triton X-100) with protease inhibitors (10 μg/ml leupeptin, 10 μg/ml pepstatin A, 10 μg/ml chymostatin) and subsequently triturated with a 25G needle. Protein G Dynabeads (ThermoFisher) were prepared according to the manufacturer’s instructions. 100 μg of lysates or 100 ng of TDP-43 recombinant protein (NM_007375, OriGene) was incubated with 30 pmol of biotin-labelled RNA for 1 h at 4°C. Proteins were eluted with 2.5× Laemmli buffer and immediately analysed by immunoblotting with rabbit anti-TDP-43 (10789-2-AP; Proteintech) and rabbit anti-hnRNP L (156682; Abcam). Probes: positive control *GRN* (nt262-288): biotin-UGUGUGUGUGUGCGCGUGUGCGUGUG; negative control (AC)12: biotin-ACACACACACACACACACACACAC; *G3BP1* (nt334-358): biotin-UUUUUUGUGUGUUAAUGGUGUGUG.

### Recombinant protein expression and purification

The RRM1 and RRM2 domains of TDP-43 (aa 102-269) in the WT and F147L/F149L mutant forms were subcloned from the pCS2 vector (Kabashi *et al*., 2010) to the pET21b vector (Novagen) modified to include a TEV cleavage site, and the resulting plasmids were transformed into *E. coli* host strain BL21 (DE3). The bacteria were grown at 37°C in Luria-Bertani (LB) media, and protein expression was induced with 1 mM isopropyl-β-D-1-galactopyranoside (IPTG) for 4 h at 30°C. The cells were harvested by centrifugation and resuspended in lysis buffer (25 mM HEPES pH 8.0, 500 mM NaCl, 1 mM DTT) supplemented with 0.15% (w/v) Protease Inhibitor Cocktail (Sigma-Aldrich). The cells were lysed by French press, sonicated for 10 sec, and centrifuged at 11,000*g* for 30 min at 4°C. The supernatant was incubated at 4°C for 1 h with Ni-charged IMAC Sepharose 6 Fast Flow (GE Healthcare). The resin was then washed three times with His-A buffer (lysis buffer + 20 mM imidazole). The bound His_6_-tagged TDP-43_102-269_ fusion protein was eluted from the washed resin by two 10-min incubations at room temperature with His-B buffer (lysis buffer + 30 mM imidazole) followed by centrifugation. The supernatant containing the protein of interest was dialysed overnight at 4°C in FPLC-A buffer (25 mM HEPES pH 8.0, 100 mM NaCl, 1 mM DTT) supplemented with TEV protease (kindly provided by J.G. Omichinski, Université de Montréal). The retentate was then applied to a Q Sepharose High-Performance column (GE Healthcare; 60-mL bed volume) equilibrated with FPLC-A buffer. The protein was eluted from the column using a gradient (from 0% to 50% over 450 mL) of FPLC-B buffer (FPLC-A + 1 M NaCl). The pooled fractions containing the protein of interest were dialyzed overnight at 4°C in storage buffer (25 mM HEPES pH 8.0, 100 mM NaCl, 2 mM DTT, and 20% glycerol) and the protein was stored at −80°C. The purity of the protein (>98%) was assessed by Coomassie-stained SDS-PAGE.

### RNA preparation

The TDP-43 binding sequence of human *G3BP1* 3’-UTR (G3BP1-RNA_32_: 5’-TTGTGTGTTAATGGTGTGTGCTCCCTCTCCA-3’) was first cloned into the pARiBo4 plasmid (Salvail-Lacoste *et al*., 2013). After plasmid linearization with EcoRI, the ARiBo-fusion RNA was *in vitro* transcribed for 3 h at 37°C using standard conditions(Salvail-Lacoste *et al*., 2013). The RNA was then purified using the ARiBo affinity purification method under non-denaturing condition (Di Tomasso *et al*., 2011), as previously described (Salvail-Lacoste *et al*., 2013). The purified RNA was concentrated with Amicon Ultra-4 centrifugal filter devices (Millipore), and exchanged in H_2_O. For radioactive labeling of the RNA, a phosphatase alkaline treatment was first performed for 1 h at 37°C, followed by inactivation of the enzyme at 65°C for 15 min. The RNA was then 5’-end-labeled with γ-(^32^P) ATP (PerkinElmer) using T4 polynucleotide kinase (New England Biolabs) according to the manufacturer’s instructions and then further purified by 15% denaturing gel electrophoresis. After gel extraction, the ^32^P-labeled RNA was resuspended in TE buffer (10 mM Tris pH 7.6, 1 mM EDTA) and stored at −20°C.

### Determination of dissociation constants (K_d_) by electrophoretic mobility shift assay (EMSA)

The ^32^P-labeled RNA was first refolded by heating 2 min at 95°C and snap-cooling on ice for 5 min. The protein samples were diluted in EMSA buffer (50 mM Tris pH 7.5, 50 mM NaCl, 0.05% NP40, 2 mM DTT, and 20% glycerol), and the concentrations were adjusted to span from 0.01X to 5X of the estimated *K_d_*. The binding reactions (20 μL) were initiated by mixing 10 pM of ^32^P-labeled RNA with the diluted protein and incubating at 4°C for 30 min. For each *K_d_* determination, binding reactions were loaded on a 10% native polyacrylamide gel (37.5:1 polyacrylamide/bisacrylamide) and run in Tris-Glycine buffer (25 mM Tris-base, 200 mM glycine) at 200 V for 2 h, with active water cooling in a cold room. The gels were then dried and exposed overnight to a storage phosphor screen. The ^32^P-labeled RNA was visualized with a Bio-Rad Personal Molecular Imager and band intensities were quantified using the ImageLab software (version 5.2.1, Bio-Rad). The fraction of bound RNA was plotted against protein concentration, and the data fitted to the Hill equation by non-linear regression analysis within the OriginPro 2015 software (OriginLab). Four independent *Kd* determination experiments were performed for the WT TDP-43_102-269_. The reported *K_d_* reflects the average values and the standard deviations from these multiple experiments.

### Sciatic axotomy and immunofluorescence

The use of animals and all procedures were performed according to the guidelines of the Canadian Council on Animal Care and were approved by the CRCHUM Institutional Committee for the Protection of Animals. Female C57BL/6 mice (6-8 weeks, ~20 g) were purchased from Charles River Laboratories (Kingston, NY, USA). We conducted our protocol based on a previously published protocol for axotomy (Moisse *et al*., 2009). Mice were weighed and anesthetized by exposure to 1.5 l/min oxygen and 4% isoflurane. After loss of limb reflexes, animals were transferred to a mask system and maintained with 1 l/min oxygen and 2% isoflurane. For a medial axotomy, mice were shaved and the site of surgery sterilized. The right sciatic nerve was exposed with an incision 1 cm below the exit from the pelvic bone. The nerve was then cut 1 cm distal from the exit point of spinal nerve roots. A surgical sterile sponge soaked in 5% fluorogold (Fluorochrome, LLC) in sterile saline was deposit at the site of the nerve cut to enable visualization of injured motor neurons post-injury. Mice were allowed to recover in a clean heated cage and allowed free access to food and water. Pain was managed with buprenorphine injection just prior to surgery, followed by a slow release formulation injected 5 h post-surgery. Neurobehavioral assessments based on a previously published scale (Rodriguez *et al*., 2006; Moisse *et al*., 2009) were conducted at days 1, 3, 5 and 7 post injury. Following transcardiac perfusion with saline and 4% paraformaldehyde, tissues were cryopreserved, and subsequently sectioned, then permeabilized and blocked with 0.5% Tween-20 and 5% normal donkey serum (Jackson ImmunoResearch, 017-000-121) for 45 min at room temperature. Primary antibody incubations were performed in 0.3% Tween-20 in PBS overnight at 4°C, followed by an appropriate fluorescently conjugated secondary antibodies against the desired species (Jackson ImmunoResearch). Antibodies used were mouse anti-TDP-43 (1:500, R&D, MAB7778), rabbit anti-G3BP1 (1:4500, Proteintech, 13057-2-AP), and guinea pig anti-FluoroGold (1:750, NM-101 FluGgp, Protos Biotech Corp.). Images were collected using a confocal microscope (SP5; Leica) equipped with LAS AF software (Leica) for acquisition at 63x. Adobe Photoshop CC 2018 was used for quantification of fluorescence intensity. A line was drawn across the longest axis of the neuron and the highest intensity along that line was recorded. This maximum intensity of a Fluorogold-positive neuron was expressed relative to the maximum intensity mean of 10 Fluorogold-negative neurons. 30 neurons (10 neurons from 3 mice each) were quantified. Importantly, all compared images were acquired with the same microscope settings.

### RNAscope and immunofluorescence on patient autopsy material

Autopsy tissues from ALS and ALS/FTLD cases were collected in accordance with the local ethics review board at Sunnybrook Health Sciences Centre, Toronto. Orbitofrontal cortex (Broadmann Areas 11 and 47) from four sporadic ALS cases with FTLD, as assessed by presence of TDP-43 pathology (ALS/FTLD-TDP-43), and four sporadic ALS cases without FTLD were used for the study (Supplementary Table II. Formalin-fixed, paraffin-embedded (FFPE) tissue from the orbitofrontal cortex of each case was trimmed and cut into 6 μm sections using a microtome. Slides were dry oven-baked for 20 min at 60°C followed by deparaffinization with fresh xylene and dehydration with sequential ethanol dilutions in water (100%, 95%, 75%, 50%). Endogenous peroxidase was inactivated by RNAscope Hydrogen Peroxide Solution (Advanced Cell Diagnostics, 322335) and antigen target retrieval was achieved by heating the samples in RNAscope Target Retrieval reagent (Advanced Cell Diagnostics, 322000) at 100°C for 30 min before the sections were pretreated with RNAscope Protease Plus (Advanced Cell Diagnostics, 322331) at 40°C for 30 min. Sections were then hybridized with Human *G3BP1* RNAscope Probes (Advanced Cell Diagnostics, 567861) at 40°C for 2 h and the resulting signals were amplified and developed with RNAscope 2.5 HD Detection Kit-Red Advanced Cell Diagnostics, 322360) according to the manufacturer’s instructions. After hybridization, sections were washed with PBS-TX (PBS, 0.15% Triton X-100) and blocked with 4% normal donkey serum in PBS-TX at RT for 1 h. Blocked sections were incubated with TDP-43 antibody (1:1000, Proteintech, 10782-2-AP) at 4°C overnight. After 3 x 5 min washes in PBST, secondary incubation was performed at ambient temperature with donkey α-rabbit 488 Alexa Fluor secondary antibody (1:500, Invitrogen, A-21206). Finally, slides were washed 3 x 5 min with PBS-TX and once with 0.1 M phosphate buffer pH 7.4 (0.08 M sodium phosphate dibasic and 0.02 M sodium phosphate monobasic) prior to coverslipping with ProLong Gold antifade reagent with 4’,6-diamidino-2-phenylindole [DAPI] (Invitrogen, P36931).

### Micrograph acquisition and quantification

Fluorescence micrographs of orbitofrontal cortex sections from ALS without FTLD (n=4) and ALSFTLD (n=4) were captured using a Leica DMI6000B microscope with a 100X objective lens on the Volocity Acquisition Suite (v6.3, Perkin Elmer). For each section, a 2-dimensional z-stack projection was produced from 25 micrographs captured at a depth of 0.25 μm through the z-axis. To allow for the quantification of *G3BP1*-positive mRNA granules (red channel) in each neuron with or without TDP-43 pathology (green channel), the green channel intensity was sufficiently enhanced to reveal cell boundaries. The number of *G3BP1*-positive mRNA granules were quantified from 25 randomly selected neurons demonstrating normal nuclear TDP-43 in the orbitofrontal cortex of ALS without FTLD (n=4) and ALS/FTLD-TDP-43 (n=4) (100 neurons total from each condition). The same sampling strategy was employed resulting in the quantification of *G3BP1*-positive mRNA granules from an additional 100 neurons demonstrating TDP-43 pathology in ALS/FTLD-TDP-43 cases (n=4). Statistical analysis comparing the mean number of *G3BP1*-positive mRNA granules in neurons with normal nuclear TDP-43 in ALS without FTLD-TDP-43 pathology and ALS/FTLD-TDP-43 versus neurons with nuclear depletion and cytoplasmic TDP-43 aggregation in ALS/FTLD-TDP-43) was performed using paired one-way ANOVA (p<0.0001).

### RNA-seq analysis

The RNA-sequencing data from human frontal cortex and cerebellum (Prudencio *et al*., 2015) (GEO accession: GSM1642314; SRA study: SRP056477) were downloaded and run in a bioinformatics pipeline using the Compute Canada clusters. Alignment was performed using HiSAT2 (Kim *et al*., 2019) against reference genome Hg38. Read counts were obtained with HTSeq-count (Anders *et al*., 2015) and differential expression analysis were performed with the Bioconductor R package DESeq2 (Love *et al*., 2014). Data were normalized using DESeq2’s median of ratios method. Polyadenylation site usage in human lumbar spinal motor neurons (Krach *et al*., 2018) (GEO accession: GSE103225; SRA study: SRP116386) was determined with QAPA (Ha *et al*., 2018).

### Statistics

Data were graphed and analyzed using Prism version 6.00 (GraphPad Software). Statistical tests are stated in the figure legends. Data were compared via two-tailed paired and unpaired t test, Mann-Whitney, one-way and two-way ANOVA with statistical significance established at p<0.05.

## RESULTS

### TDP-43 nuclear depletion and disease-associated TDP-43 variants decrease G3BP1 levels

We have previously demonstrated that depletion of TDP-43 results in a downregulation of G3BP1 protein levels (McDonald *et al*., 2011). As ALS/FTLD cases predominantly display a cytoplasmic mislocalization of TDP-43 concomitant with its nuclear depletion, we investigated whether cytoplasmic-restricted TDP-43 could rescue G3BP1 protein levels. To this end, we expressed an siRNA-resistant cDNA encoding TDP-43 with an inactivated nuclear localization signal (FLAG-TDP-43^δNLS^) following siRNA-mediated depletion of endogenous TDP-43 in HeLa cells. Immunofluorescence analysis confirmed the cytoplasmic expression of the exogenous construct FLAG-TDP-43^δNLS^ in comparison to mock transfected control cells (**Fig. S1A, B**), as previously reported (Winton *et al*., 2008). Expression of FLAG-tagged TDP-43^ΔNLS^ in the context of endogenous TDP-43 depletion yielded a ~50% decrease in G3BP1 protein levels (siCTL+mock vs siTDP-43+TDP-43^ΔNLS^, p=0.0029) comparable to G3BP1 levels in TDP-43 siRNA treated cells co-transfected with empty plasmid (**Fig. 1A, B)**. Thus, cytoplasmic-restricted TDP-43 does not rescue G3BP1 protein levels.

**Figure 1.**
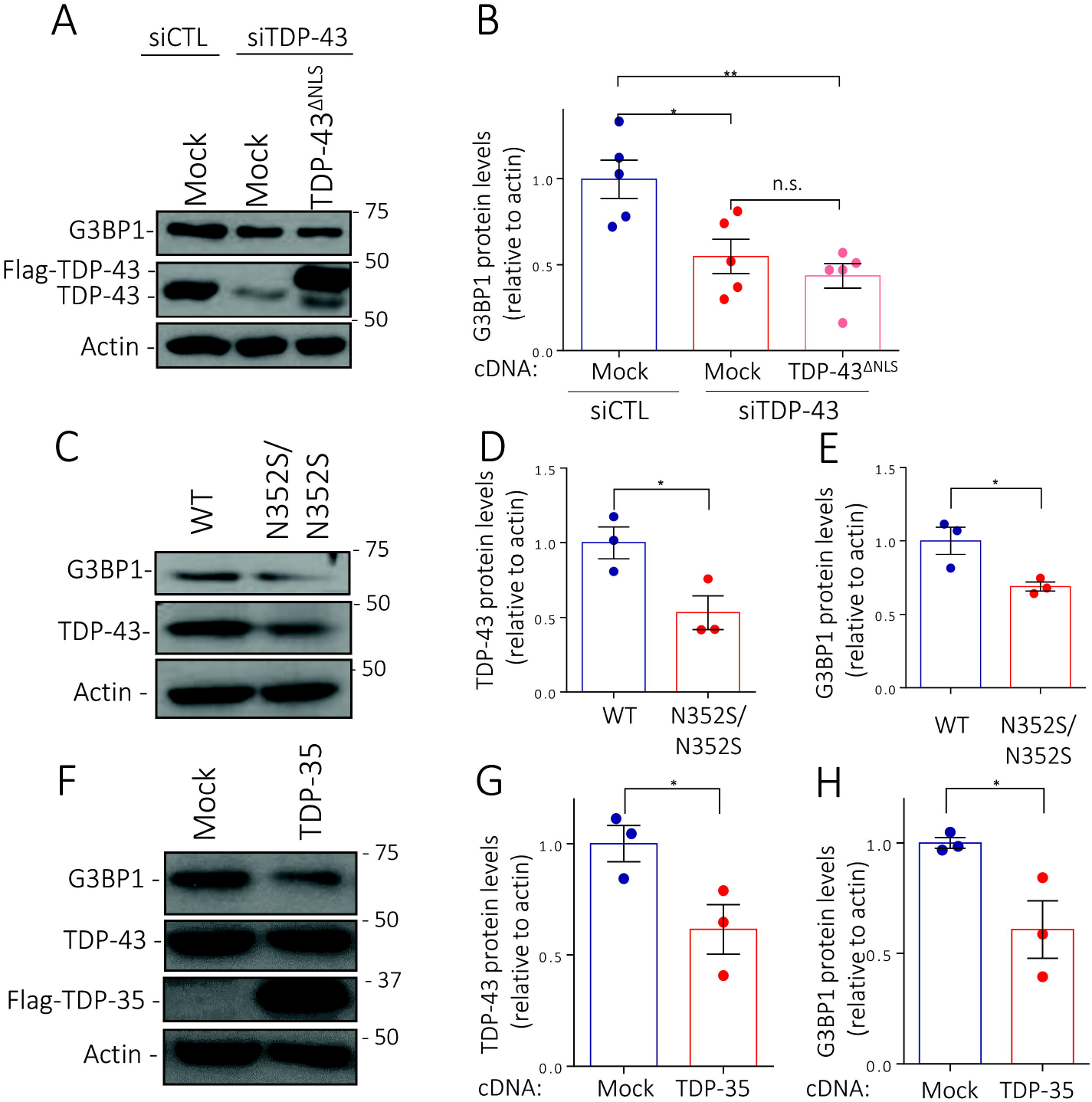
TDP-43 nuclear depletion and ALS-related species induce G3BP1 down-regulation. (A) HeLa cells were transfected with siTDP-43 or siControl (siCTL), then transfected with empty plasmid (Mock) or Flag-TDP-43^ΔNLS^ and immunoblotted. (B) Quantification via densitometry of G3BP1 protein levels normalized to actin. (C) Protein from SH-SY5Y WT and TDP-43^N352S/N352S^ cells were extracted, and immunoblotted. Quantification via densitometry of (D) TDP-43 and (E) G3BP1 protein levels normalized to actin. (F) HeLa cells were transfected with empty plasmid (Mock) or Flag-tagged TDP-35 and immunoblotted. Quantification via densitometry of (G) TDP-43 and (H) G3BP1 protein levels normalized to actin. Data from 3-5 independent experiments are expressed as the mean fold change ± SEM; Unpaired t test \**p*<0.05; \*\**p*<0.01.

Overexpression studies of TDP-43 ALS-associated mutations have reported defects in stress granule disassembly and/or cytoplasmic aggregation (Liu-Yesucevitz *et al*., 2010; Dewey *et al*., 2011). However, this is a phenotype frequently observed in studies involving the overexpression of aggregation prone RBPs. Thus, we opted to evaluate the impact of a familial ALS-causing mutation (N352S) genome edited onto both endogenous TDP-43 alleles in SH-SY5Y cells (Melamed *et al*., 2019). Consistent with Melamed et al., we noted a 30% decrease in TDP-43 protein levels (p=0.0334) which correlated with a 47% reduction in G3BP1 protein levels (**Fig. 1C, D**; p=0.0393). This reduction was also recapitulated at the mRNA level (**Fig. S1D**; p=0.0413). In addition, expression of TDP-35, an N-terminal truncated TDP-43 variant generated via alternative transcriptional start codon usage that is used more often in ALS patients and is predominantly localized into cytoplasmic inclusions (Xiao *et al*., 2015), also compromised G3BP1 protein levels (**Fig. 1E, F, S1C**; p=0.0065). Here, we also noted that endogenous TDP-43 levels were reduced 40% (p=0.0489). Thus, expression of two pathological TDP-43 variants, which are associated with decreased endogenous TDP-43 levels, resulted in reduced G3BP1 protein.

### TDP-43 uniquely modulates one of two possible *G3BP1* transcripts

According to the Genotype-Tissue Expression (GTEx) project, two transcripts encode the same G3BP1 protein (ENT00000394123.7 and ENST00000356245.7), the latter of which has only recently been annotated. The two transcripts primarily differ in the size of their 3’UTRs (8,901nt vs. 1,466nt), with the first 1,466nt being shared between the two isoforms (**Fig. 2A**). Using primer pairs to the distal 3’ end of the longer transcript, we confirmed the presence of the longer 3’UTR-containing transcript to varying amounts in five different cell lines via RT-PCR (**Fig. 2B, C**). Our previous work indicated that siRNA-mediated depletion of TDP-43 decreased *G3BP1* mRNA levels (McDonald *et al*., 2011), but these studies were performed using probe sets targeting exonic *G3BP1* sequence, and therefore were unable to differentiate between long and short *G3BP1* isoforms. To address the question of whether TDP-43 regulates one or both transcripts, we knocked down TDP-43 in several cell lines and separately assessed total and long *G3BP1* isoforms. Downregulation of TDP-43 using siRNA in HeLa, SK-N-SH and SH-SY5Y lowered total *G3BP1* levels (HeLa: p=0.0222; SK-N-SH: p=0.0067; SH-SY5Y: p=0.0271, consistent with our previous results (McDonald *et al*., 2011; Aulas *et al*., 2012) (**Fig. 2D**). However, using a probe set to uniquely identify *G3BP1* long transcripts, we found that the longer isoform is unaffected by TDP-43 knockdown in all three cell types tested (**Fig. 2E-F**). This suggests that it is the shorter *G3BP1* transcript that is modulated by TDP-43.

**Figure 2.**
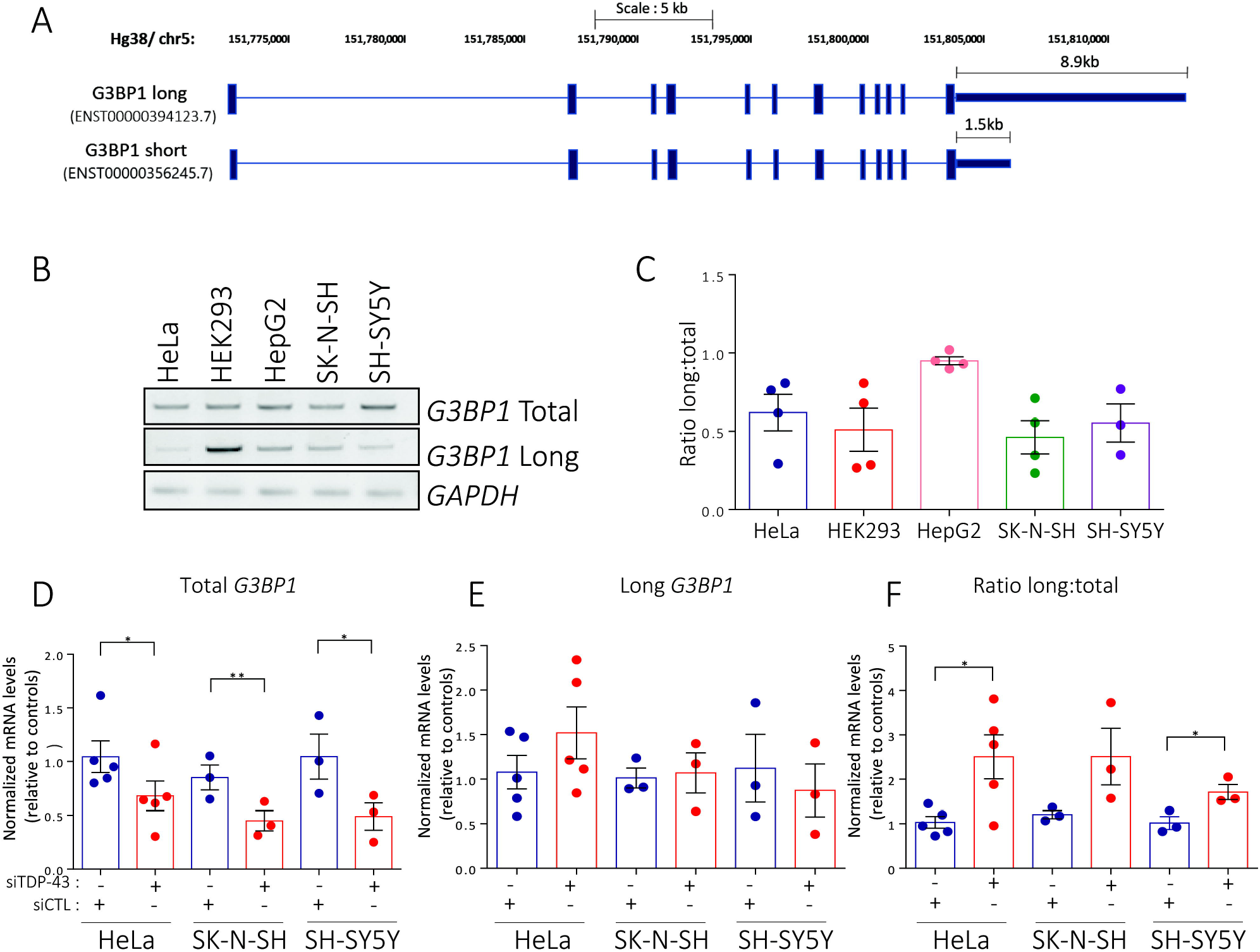
TDP-43 depletion only affects *G3BP1* short transcript. (A) Schematic of Genome Reference Consortium Human GRCh38.p12 *G3BP1* transcripts. Long transcript has 8,901nt 3’UTR while the shorter version has a 1,466nt 3’UTR. (B) RT-PCR for total *G3BP1*, long *G3BP1* transcripts and *GAPDH* of extracted RNA from HeLa, HEK293, HepG2, SK-N-SH and SH-SY5Y cells, n=3-4. (C) Quantification via densitometry of the ratio long:total transcripts and normalized to *GAPDH*, n=3-4. (D-E) qRT-PCR for total *G3BP1* and long *G3BP1* transcripts extracted RNA from HeLa, SK-N-SH and SH-SY5Y cells transfected with siCTL or siTDP-43 and normalized with *GAPDH* and *18S*. (F) Ratio of the levels of long *G3BP1* transcripts:total *G3BP1*, mean ± SEM, n=3-5, Unpaired t test *p<0.05.

### TDP-43 stabilizes the short *G3BP1* isoform via its 3’UTR

Since TDP-43 depletion is associated with a downregulation of the short *G3BP1* transcript, we first sought to determine whether this was mediated by the promotor or the 3’UTR. Using a reporter assay in which the promotor and short 3’UTR were each fused to an optimized *Renilla* reporter gene (RenSP), siRNA-mediated depletion of TDP-43 reduced steady state luciferase activity of the *G3BP1* 3’UTR construct by 43% compared to control (*GAPDH*, **Fig. 3A**, p=0.042). In contrast, the luciferase activity of the *G3BP1* promotor reporter was not significantly changed compared to controls (**Fig. S2**). mRNA abundance is frequently governed by 3’UTR elements linked to mRNA stability. To evaluate whether TDP-43 stabilizes *G3BP1* mRNA, we employed a doxycycline-repressible transcript containing His-tagged G3BP1 cDNA fused to the short *G3BP1* 3’UTR. This approach allowed us to specifically determine exogenous *G3BP1* mRNA decay without altering total cellular metabolism (as is observed with Actinomycin D treatment) (Baird and Hogg, 2018). In TDP-43-depleted conditions, the exogenous *G3BP1* transcript was four-fold less stable than in siControl treated cells (t_1/2_ siCTL: 4.3 ± 1.0 h vs. t_1/2_ siTDP-43: 1.2 ± 0.4 h, p=0.0179; **Fig. 3B, C**). Thus, TDP-43 is required to stabilize the short *G3BP1* transcript.

**Figure 3.**
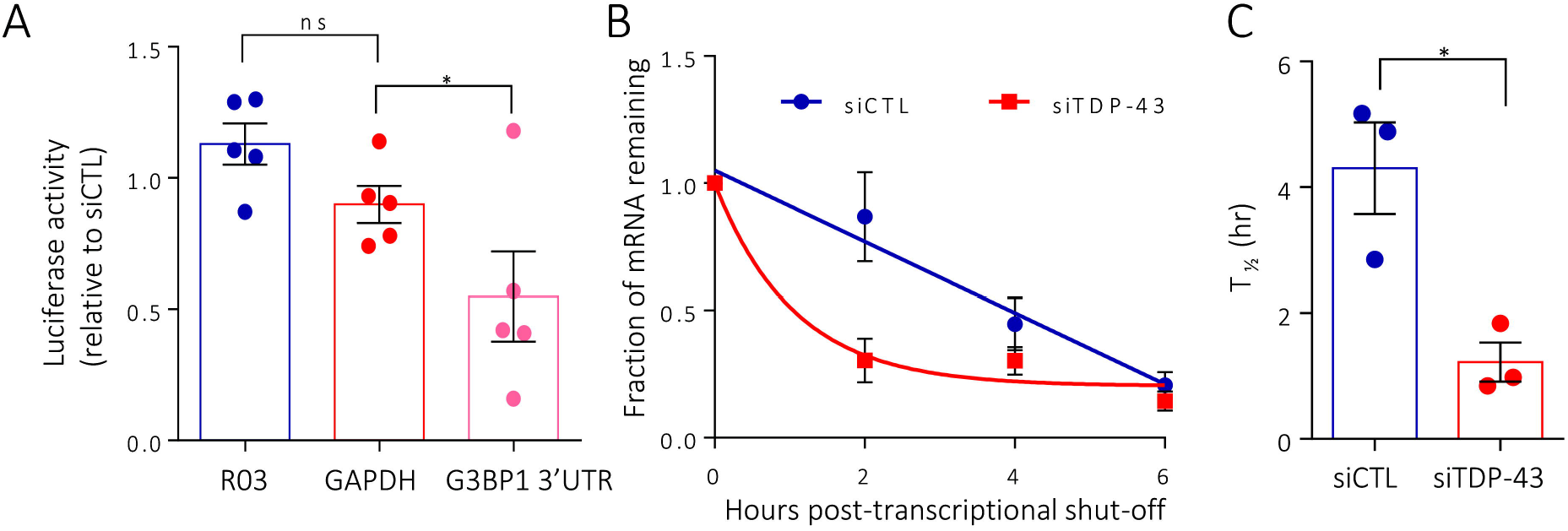
TDP-43 regulates and stabilizes *G3BP1* via its 3’UTR. (A) HeLa cells were transfected with siTDP-43 or siCTL, then co-transfected with the indicated reporter plasmids. Luciferase activity of *G3BP1* 3’UTR is expressed relative to siCTL cells. *GADPH* and R03 (random sequences) are used as controls, mean ± SEM, n=4, Unpaired t test *p<0.05. (B) qRT-PCR for *His-G3BP1* transcripts, normalized to *18S* following doxycycline treatment for 6 h to shut off expression, n=3. (C) Half-life of *His-G3BP1* transcript in HeLa-Tet-off cells transfected with siCTL compared to cells transfected with siTDP-43, n=3, Unpaired t test *p<0.05.

### The 3’UTR of *G3BP1* harbors a highly conserved UG-rich regulatory element

TDP-43 preferentially binds UG-rich sequences (Lukavsky *et al*., 2013; Mompean *et al*., 2016), which are often associated with mRNA stability (Fiesel *et al*., 2010; Gu *et al*., 2017). In order to better characterize TDP-43-dependent stabilization of *G3BP1*, we analyzed the human short 3’UTR using RBPmap, which predicts RBP binding sites (Paz *et al*., 2014). A cluster of 13 potential TDP-43 binding sites were identified, located within nt334-358 of the 3’UTR of both transcripts (GRCh38/hg38 chr5: 151,804,220-151,804,244; **Fig. 4A**). Nucleotide alignment of this sequence across 13 different species indicated an element at 3’UTR^nt341-357^ (GRCh38/hg38 chr5: 151,804,227-151,804,243) that is highly conserved across all species examined despite overall degeneration of the 3’UTR between species (**Suppl. Table I**). MEME analysis (Bailey *et al*., 2009) confirms the conservation of UG-repeats in this 16-nt element (**Fig. 4B**). TDP-43 and its orthologs are structurally conserved throughout evolution, especially within the RBD (Lukavsky *et al*., 2013). Moreover, G3BP1 is also conserved in *C. elegans*, with only one protein-encoding transcript, corresponding to the shorter isoform. Consistent with our data in human cell lines, *tdp-1* null worms (Vaccaro *et al*., 2012) exhibit a 4-fold decrease of *gtbp-1 (G3BP1* orthologue) mRNA compared to N2 control worms (**Fig. 4C, D**; p=0.0326). Thus, *G3BP1* mRNA contains a highly conserved UG-rich *cis* regulatory element in its 3’UTR that is functionally regulated by TDP-43.

**Figure 4.**
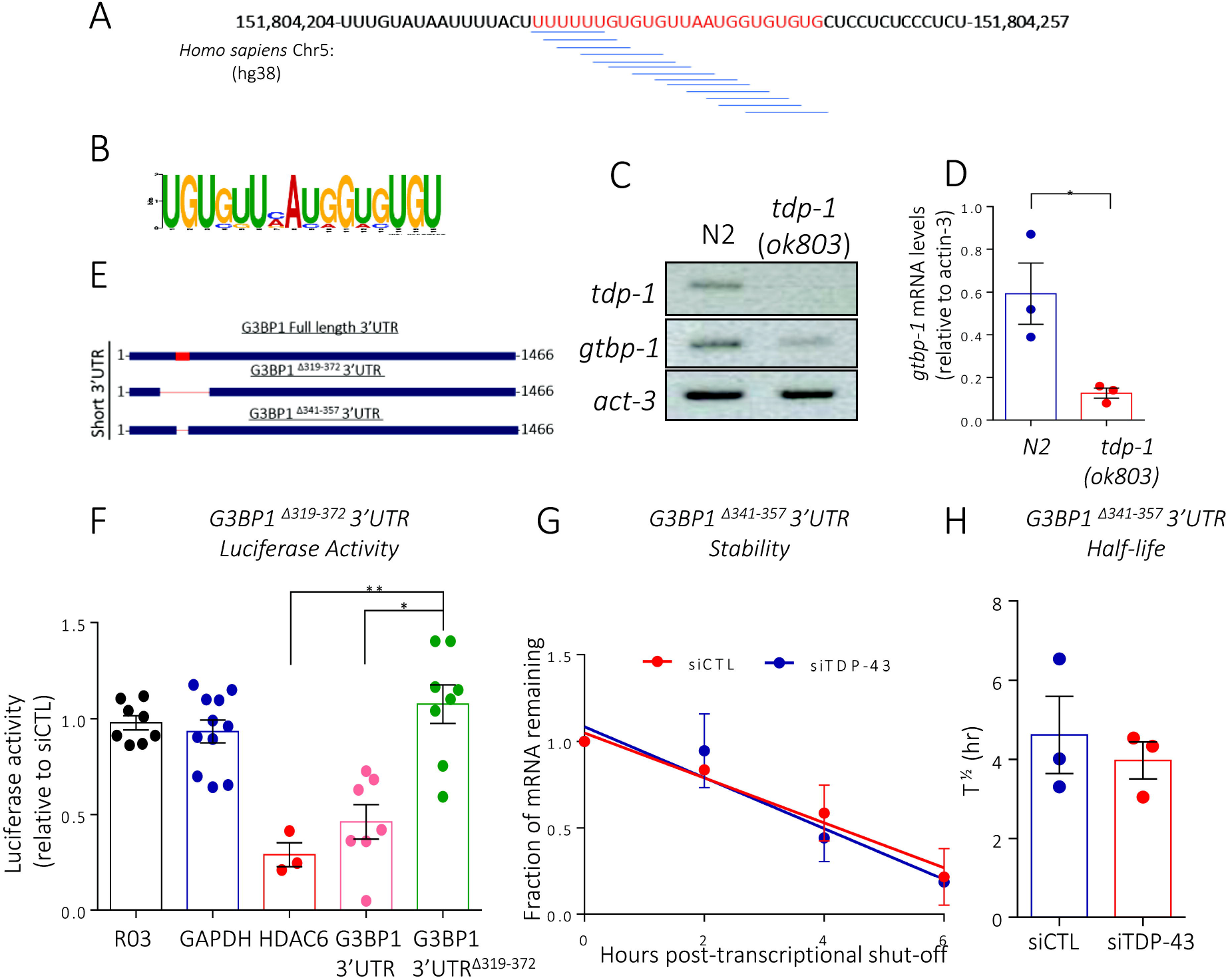
Identification of a conserved element in the G3BP1 3’UTR. (A) Schematic representation of the conserved element identified in the human *G3BP1* 3’UTR between nt319-372 (GRCh38/hg38 chr5: 151,183,766-151,183,816). The blue lines represent TDP-43 binding sites suggested by RNAmap. (B) Consensus sequence of the conserved sequence obtained from 13 species using the consensus sequence generator MEME. (C) RT-PCR for *tdp-1, gtbp-1* and *act-3* of extracted RNA from N2 (Wild-type control) and *tdp-1* null worms. (D) Quantification via densitometry of the absolute *gtbp-1* levels normalized to *act-3*. n=3, Unpaired t test *p<0.05. (E) Schematic representation of the *G3BP1* 3’UTR constructs used. The red rectangle represents the conserved element. (F) HeLa cells were transfected with siTDP-43 or siCTL, then co-transfected with the indicated reporter plasmids. *GADPH* and R03 (random sequences) are negative controls and *HDAC6* is a positive control, mean ± SEM, n=3-8, Unpaired t test *p<0.05; **p<0.01. (G) qRT-PCR for *His-G3BP1^Δ341-357^* transcript, normalized to *18S* following doxycycline treatment for 6 hours to shut off expression. (H) Half-life of *His-G3BP1^Δ341-357^* transcript in HeLa-Tet-off cells transfected with siCTL compared to cells transfected with siTDP-43, n=3, Unpaired t test.

### TDP-43 stabilizes *G3BP1* mRNA via a conserved 3’UTR regulatory element

To investigate the relevance of the identified conserved sequence, we revisited the luciferase reporter assay using a construct where a region including the regulatory element was deleted (*G3BP1* 3’UTR^Δ319-372^; **Fig. 4E**). Unlike the intact short *G3BP1* 3’UTR, or the positive control *HDAC6*, siRNA-mediated depletion of TDP-43 had no effect on the luciferase activity of *G3BP1* 3’UTR^Δ319-372^ (**Fig. 4F**). Moreover, using a doxycycline-repressible version of this construct with only the 16-nt conserved element deleted (*G3BP1* 3’UTR^Δ341-357,^ **Fig. 4E**), we determined that the half-life of this exogenous transcript was comparable in the presence and absence of TDP-43 (4.6 ± 1.7 h vs 3.9 ± 0.8 h, p = 0.58; **Fig. 4G, H**). The data obtained here indicate that the identified 16-nt UG-rich regulatory element is required for TDP-43-mediated stabilization of the short *G3BP1* transcript.

### TDP-43 directly binds the conserved element with high affinity

To determine if TDP-43 mediates stabilization of *G3BP1* mRNA via binding to the transcript, we performed immunoprecipitation of endogenous TDP-43 followed by RT-PCR of associated mRNAs. *G3BP1* transcripts were recovered from TDP-43 immunoprecipitates, as was the positive control *CAMKII* (Wang *et al*., 2008). *HSPA1A* served as a negative control (Wang *et al*., 2008) (**Fig. 5A**). Next, in order to determine if the stabilizing effect of TDP-43 on the short *G3BP1* transcript was due to binding to the identified conserved element, we performed an RNA pulldown. Using a biotinylated probe (*G3BP1* 3’UTR^nt319-372^) and HeLa whole cell extracts, we retrieved TDP-43 from the lysate, similar to a probe derived from *GRN* which has been previously reported to be bound by TDP-43 (Colombrita *et al*., 2012) (**Fig. 4B**). The binding was considered specific since an AC-rich probe [(*AC*)_12_] was not bound by endogenous TDP-43 but was by hnRNP L, an RBP with a preference for AC-rich sequences (Hui *et al*., 2005). To determine if the interaction between the 3’UTR regulatory element of *G3BP1* and TDP-43 was direct, the RNA pulldown was repeated using commercially-produced recombinant TDP-43 protein. Both the *G3BP1* and *GRN* probes, but not the (*AC*)_12_ probe, were bound by recombinant TDP-43 (rTDP-43) (**Fig. 5B**). Thus, TDP-43 directly binds the region containing the conserved regulatory element in the 3’UTR of *G3BP1*.

**Figure 5.**
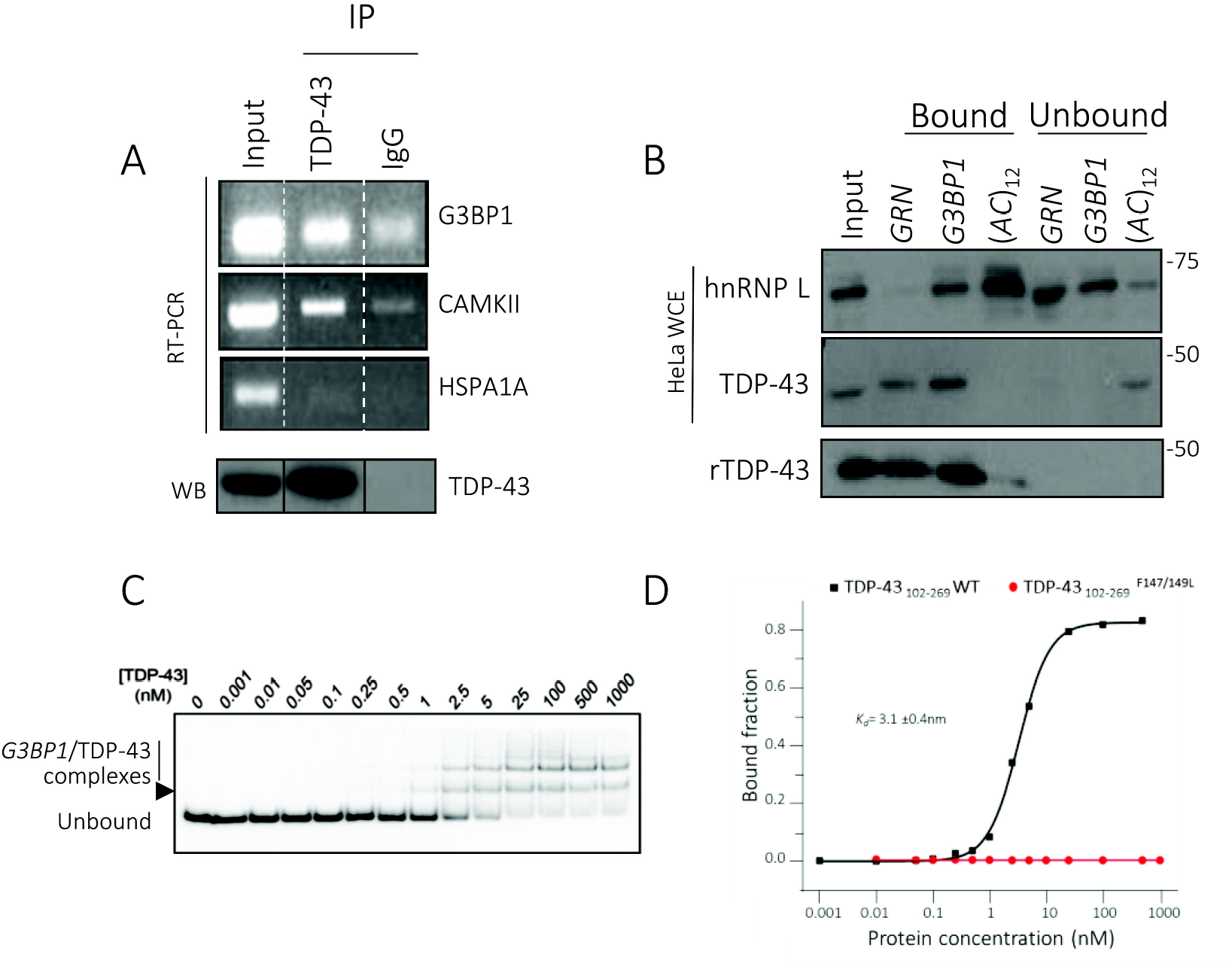
TDP-43 directly binds the conserved element. (A) TDP-43 protein and its associated transcripts were co-immunoprecipitated from HeLa cell extracts. mRNAs binding to TDP-43 were extracted, reverse transcribed and amplified for *G3BP1, CaMKII* (positive control) and *HSPA1A* (negative control), showing that TDP-43 binds *G3BP1* mRNA. Flag-IP serves as a control to demonstrate specificity of TDP-43 immunoprecipitation. Black lines indicated spliced lanes. (B) Proteins from whole HeLa cell extracts or TDP-43 recombinant protein (rTDP-43) were pulled down using biotinylated RNA probes containing the TDP-43 binding sequence in *GRN* (positive control), *G3BP1* 3’UTR^nt319-372^, or (*AC*)_12_ (negative control). hnRNP L served as a positive control for AC-repeat binding. (C) Typical EMSA performed with 10 pM 5’-[^32^P]-labeled RNA and increasing concentrations of TDP-43_102-269_. (D) Typical binding curve using WT TDP-43_102-269_ (black line) and the F147L/F149L mutant of TDP-43_102-269_ (red line).

To establish that the binding was via the RBD of TDP-43 and to determine the affinity of the interaction of TDP-43 for the 3’UTR element, we performed an *in vitro* electrophoretic mobility shift assay (EMSA) using highly-purified components. The RBD of TDP-43 (TDP-43_102-269_) efficiently bound a 32-nt RNA encompassing the *G3BP1* 3’UTR conserved regulatory sequence (G3BP1-RNA_32_) and corresponding to *G3BP1* 3’UTR^nt340-378^). This binding was accompanied by formation of two main shifted bands on the gel, as observed in previous binding studies with (TG)_12_ DNA oligomers (Buratti and Baralle, 2001). Given that the G3BP1-RNA_32_ contains two GUGUGU sequences that have the potential to serve as independent binding sites, we speculate that these two bands represent the formation of RNA-protein complexes containing either one or two molecules of TDP-43_102_-2_69_. Interestingly, the wild-type TDP-43_102_-2_69_ binds with very high affinity to G3BP1-RNA_32_. In contrast, there is no evidence of RNA binding for the TDP-43 102-269 protein fragment containing mutations that inactivate the RNA binding pocket (F147L/F149L) at concentrations up to 1 μM (**Fig. 5C, D)**. EMSA experiments performed in multiple replicates with the wild-type TDP-43_102_-2_69_ indicate an average *K_d_* value of 3.1 ± 0.4 nM. Thus, taken together, these data demonstrate a high affinity of TDP-43 for the identified conserved element in the short 3’UTR of *G3BP1*.

### Injury-induced TDP-43 nuclear efflux correlates with reduced G3BP1 in motor neurons *in vivo*

TDP-43 nuclear efflux has been reported in motor neurons of axotomized mice (Urushitani *et al*., 2007; Moisse *et al*., 2009). Thus, as a way to evaluate whether TDP-43 nuclear depletion can impact G3BP1 *in vivo*, we axotomized C57Bl/6 mice by severing the sciatic nerve just past the sciatic notch, as it exits the pelvic bone (**Fig. 6A**). The behavioural evaluation of all injured mice was performed using the previously published NBA scoring system (Rodriguez *et al*., 2006; Moisse *et al*., 2009). At day 1, axotomized mice exhibited notable paralysis of the foot with dragging, knuckle walking, and no toe extension on the injured side (**Fig. 6B**). This phenotype attenuated in the following days post-injury as shown by the significant decrease in the NBA score at day 7 post-injury (**Fig. 6B**). We confirmed the presence of axotomized motor neurons on the ipsilateral, but not the contralateral, side of the ventral spinal cord using FluoroGold retrograde labelling (**Fig. 6C**). Investigation of TDP-43 expression/localization in injured neurons labelled with FluoroGold revealed an 80% depletion of nuclear TDP-43 at day 7 post-injury. This was accompanied by a 70% reduction in G3BP1 expression in these motor neurons compared to the contralateral side (**Fig. 6C, D;** p<0.0001). To verify the specificity of these observations, we also immunolabelled for FUS (**Fig. S3**). After injury, FUS remained localized to the nucleus even at day 7, consistent with previous work indicating that FUS is not regulated by TDP-43 (McDonald *et al*., 2011; Aulas *et al*., 2012). Taken together, these results demonstrate an *in vivo* correlation between nuclear TDP-43 depletion and G3BP1 reduction in adult motor neurons.

**Figure 6.**
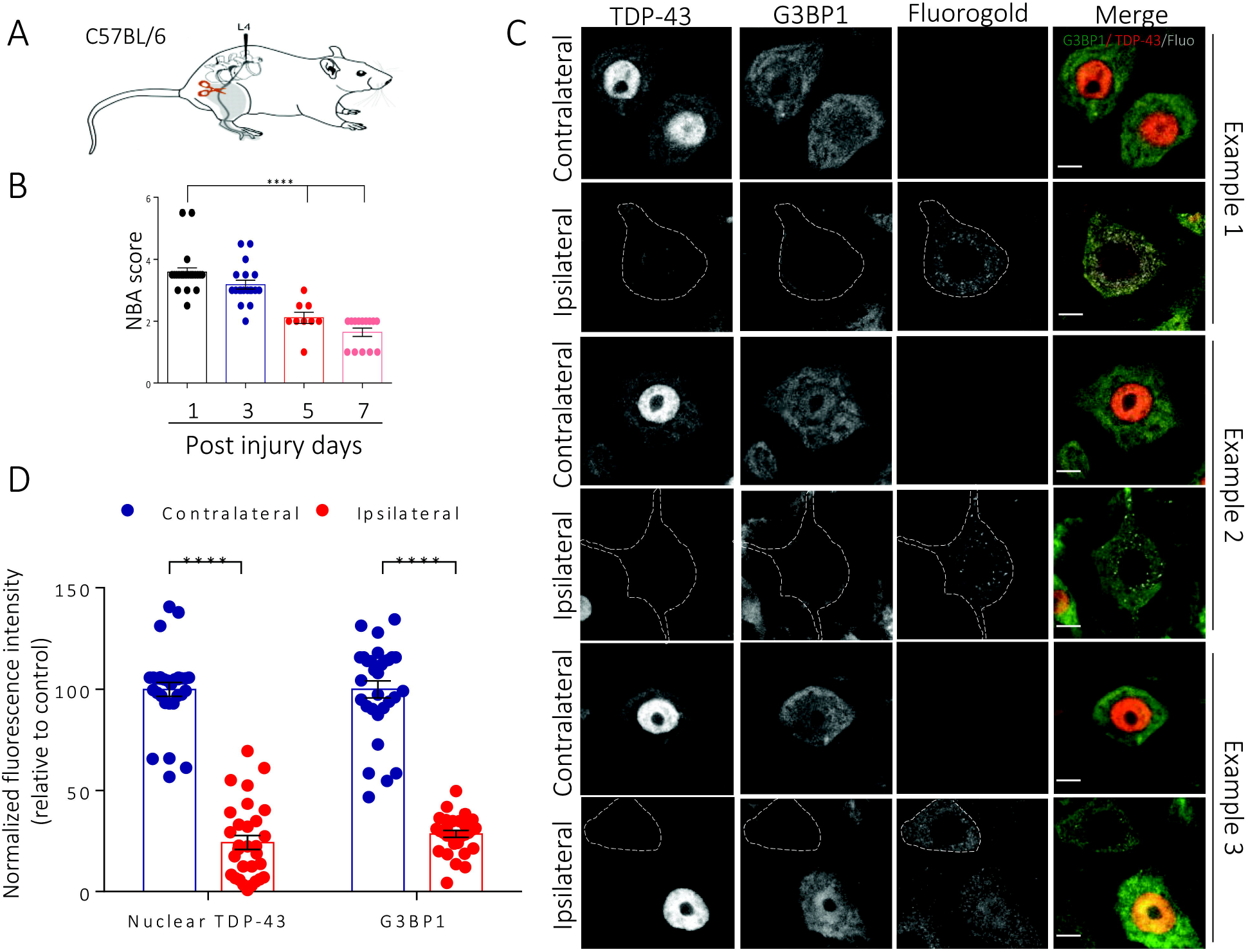
Changes in TDP-43 and G3BP1 expression in injured motor neurons following medial axotomy. (A) Schematic of the medial site of axotomy. (B) NBA score of the axotomized mice at days 1, 3, 5 and 7. Data is expressed as the mean± SEM; Unpaired t test ****p<0.0001. (C) Three representative examples of control non-axotomized (ipsilateral) and axotomized spinal neurons (contralateral) at day 7, showing injured spinal motor neurons of the ventral horn labelled for TDP-43 (red), G3BP1 (green), and Fluorogold (white). The dotted line shows the cell boundary. (D) Fluorescence intensity of nuclear TDP-43 and total G3BP1 for the control non-injured and injured motor neurons at day 7, N=3 mice, n=10 neurons per mouse. Data is expressed as the mean ± SEM; Unpaired T-test ****p<0.0001. Scale bar, 10 μm.

### The short *G3BP1* transcript predominates in affected cells types and is reduced in ALS and ALS/FTD neurons

Given that only one of the two *G3BP1* transcripts is sensitive to TDP-43 levels, it was imperative to determine the relative abundance of each transcript in cells/tissue of relevance to disease. Analysis of published transcriptomics data from two different brain regions of healthy individuals (Prudencio *et al*., 2015) revealed a paucity of reads corresponding to the extended 3’UTR region of the long *G3BP1* transcript in the frontal cortex, compared to the cerebellum (**Fig. 7A**), suggesting that the short *G3BP1* transcript predominates in the frontal cortex. Analysis of global expression of *G3BP1* (i.e. both transcripts) revealed a modest reduction in *G3BP1* expression in the frontal cortex compared to the cerebellum, which demonstrated high variability across individuals (**Fig. 7B;** log2 FC: 0.30; p = 0.04). We also examined polyadenylation site usage in published transcriptomics data generated from lumbar spinal motor neurons isolated by laser capture microdissection from healthy individuals (Krach *et al*., 2018). This analysis clearly indicated the preferential utilization of the proximal polyadenylation site, reflecting that the short *G3BP1* transcript is more abundant than the longer isoform in the most ALS-vulnerable neurons (**Fig. 7C**; p=0.0004). Both of these aforementioned datasets also included ALS and/or ALS/FTD cases. However, we were unable to detect a significant difference between *G3BP1* 3’UTR usage between controls and individuals with ALS or ALS/FTD, presumably due to the relatively low overall abundance of *G3BP1* transcripts (data not shown). Thus, we assessed *G3BP1* transcript levels in human neurons differentiated from induced pluripotent stem cells (iPSCs) via Ngn1-2 expression to yield a near-homogenous population of iPSC-derived neurons with properties consistent with glutamatergic, excitatory forebrain-like neurons (i^3^Neurons) (Zhang *et al*., 2013; Busskamp *et al*., 2014; Lam *et al*., 2017b). These i^3^Neurons were modified to express a single copy of wild type or mutant (M337V) TDP-43. Via qRT-PCR, we observed that *G3BP1* total transcript levels, but not the long transcript, were reduced 49% in TDP-43^M337V^ expressing i^3^Neurons compared to TDP-43^WT^ i^3^Neurons (**Fig. 7D**, p=0.0272). In agreement with our data, this result suggests that it is the shorter *G3BP1* transcript impacted by the TDP-43 mutation.

**Figure 7.**
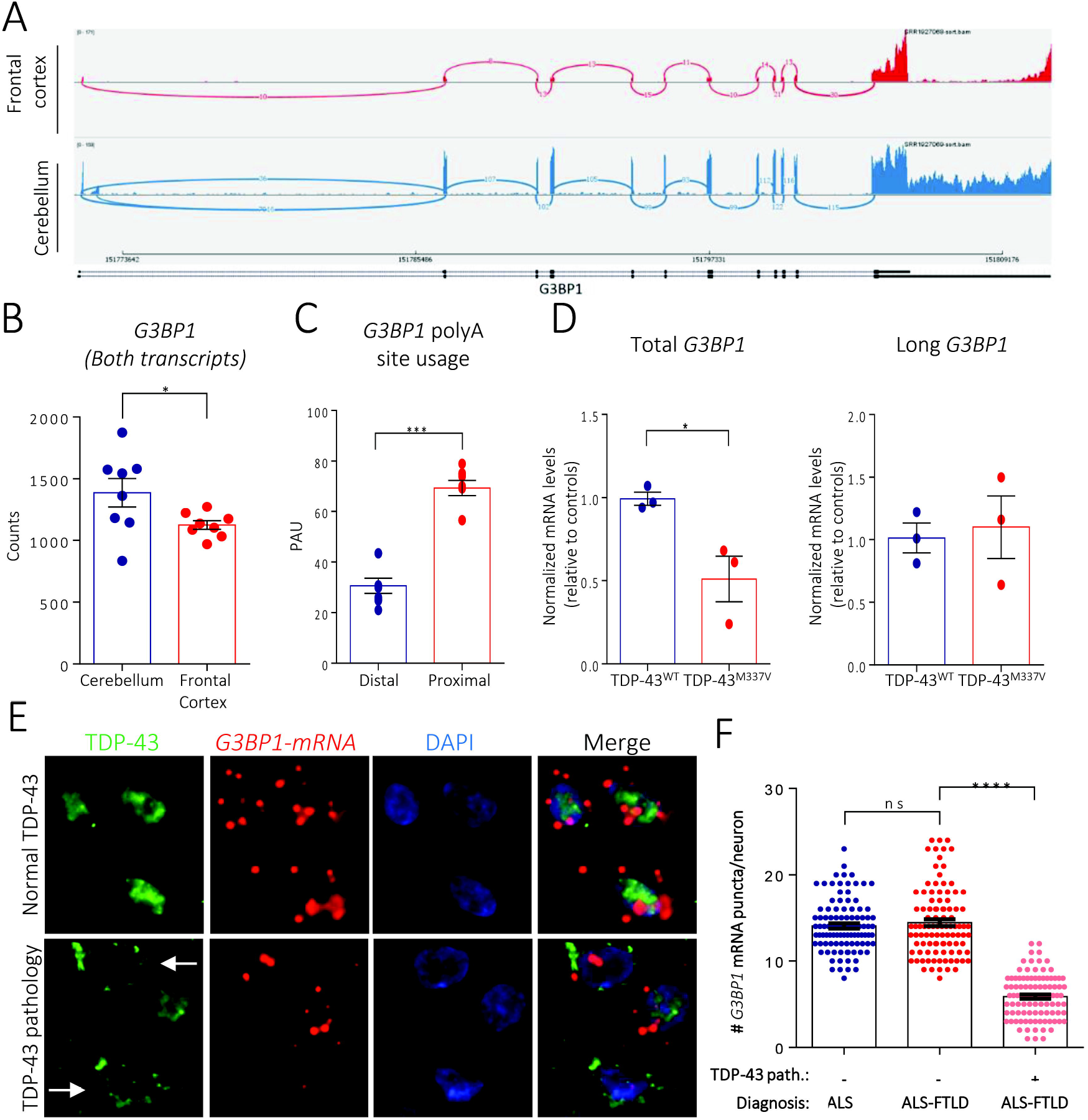
*G3BP1* short mRNA is destabilized and reduced in patients with TDP-43-mediated neurodegeneration. (A) Sashimi plots of GEO dataset GSE67196 for human *G3BP1* in cerebellum and frontal cortex of control case, showing reduced reads for the long G3BP1 3’UTR in the frontal cortex. (B) Comparison of *G3BP1* levels (both transcripts) in human cerebellum and frontal cortex in healthy controls. *p<0.05. (C) Quantification of polyadenylation site usage in *G3BP1* transcripts in lumbar spinal motor neurons isolated by laser capture microdissection (GEO dataset GSE103225). Data is expressed as the mean± SEM; Unpaired t test ***p<0.001. (D) qRT-PCR for total and long *G3BP1* transcripts extracted RNA from i3Neurons expressing a single inserted copy of wild type and mutant TDP-43. Data is normalized to *GAPDH* and *18S* and expressed as the mean ± SEM, n=3, Paired t test *p<0.05. (E) Neurons in the orbitofrontal cortex of ALS and ALS/FTLD cases showing normal nuclear localization of TDP-43 (top row, green) or nuclear TDP-43 depletion/cytoplasmic TDP-43 accumulation (bottom row, green). *G3BP1* mRNA (red dots) is labelled with RNAscope probes, nuclei are marked with DAPI (blue). Arrows show neurons with TDP-43 pathology. (F) Quantification of *G3BP1* mRNA signals in neurons of the orbitofrontal cortex of ALS and ALS/FTLD with or without TDP-43 pathology (defined as obvious cytoplasmic accumulations or reduced nuclear TDP-43 levels). Data is expressed as the mean± SEM; One-way Anova ****p<0.0001. Scale bar, 10μm.

Finally, to directly quantify total *G3BP1* transcripts in the context of TDP-43 pathology, we performed quantitative *in* situ hybridization using RNAscope probes to *G3BP1* coupled with immunofluorescence labeling using TDP-43 antibody on orbitofrontal cortices of ALS/FTLD cases with cortical TDP-43 pathology in comparison with ALS cases without FTLD and no cortical TDP-43 pathology (n=4 each, **Suppl. Table II**). The number of *G3BP1* transcripts detected using RNAscope was equivalent in cortical neurons with normal nuclear TDP-43 localization (**Fig. 7E**) in both ALS/FTLD-TDP-43 cases and ALS/no FTLD cases (14.1 ± 3 and 14.4 ± 4, respectively, n=100 neurons per group, **Fig. 7F**). In contrast, *G3BP1* transcript levels were reduced 60% in ALS/FTLD neurons with TDP-43 pathology (5.9 ± 2.5 puncta per neuron; n=100 neurons, p=2.5 ×10^-35^; **Fig. 7E, F**), as evidenced by nuclear depletion and cytoplasmic mislocalization of TDP-43 (**Fig. 7E**, indicated with arrows). Taken together, these data are consistent with the concept that *G3BP1* mRNA is destabilized in the absence of nuclear TDP-43 in disease affected neurons in ALS/FTLD.

## DISCUSSION

In this study, we demonstrate the mechanism by which the loss of TDP-43 leads to reduced levels of *G3BP1* mRNA and protein (McDonald *et al*., 2011; Aulas *et al*., 2012; Aulas *et al*., 2015). We found that TDP-43 binds to and regulates the *G3BP1* short transcript via its 3’UTR, similar to that described for *HDAC6, ADD2* and *MAPT*(Fiesel *et al*., 2010; Colombrita *et al*., 2012; Costessi *et al*., 2014; Gu *et al*., 2017). TDP-43 depletion reduces the stability of the most abundant *G3BP1* transcript, explaining the observed decrease in G3BP1 protein level in TDP-43-depleted cells. In general, cellular stress and pathology are associated with an increase in the proportion of shorter transcripts, suggesting that this TDP-43 regulated transcript may become elevated in adverse conditions (Lau *et al*., 2010; Romo *et al*., 2017; Zheng *et al*., 2018) and thus outcompete the longer transcript for translation. Moreover, short and long 3’UTRs can be differentially regulated with long 3’UTR transcripts being longer-lived (Tushev *et al*., 2018). A number of factors, including RNA structure and competitive/collaborative binding with other RBPs or miRNAs can impact translatability and localization of a given transcript. A well-known example is *BDNF*, where the isoform with the short 3’UTR is restricted to the perinuclear area while the longer transcript is more broadly distributed in the soma (Lau *et al*., 2010). It remains to be determined if this is relevant to *G3BP1* and why it is impervious to TDP-43 regulation.

TDP-43 stabilizes *G3BP1* transcripts via an evolutionary conserved *cis* regulatory sequence located within the 3’UTR. In alignment with recent findings of the ENCODE project, the identified regulatory element is UG-rich but not a pure UG cluster (Van Nostrand *et al*., 2020). TDP-43 directly binds the regulatory element with a *K_d_* that is 6-23x higher than that reported for small UG-rich RNAs (20-70 nM) (Lukavsky *et al*., 2013), but is similar to that reported for a 16-nt (2.8 nM) (Kuo *et al*., 2014) and a 30-nt RNA (5 nM) (Kuo *et al*., 2014) consisting entirely of UG repeats. While the entire *G3BP1* 3’UTR sequence is poorly conserved amongst the 13 species examined (and 10 additional species, data not shown), the 16-nt element we identified is very highly conserved and functional, as supported by our data in *C. elegans* where the sequence is 67% conserved. Together, these findings indicate that the regulatory element is necessary and sufficient for TDP-43 mediated stabilization of *G3BP1* mRNA. Interestingly, the conserved element we have identified lies within a region that harbors very few SNPs (UCSC genome browser, VarSome), which demonstrates a conservation of the sequence and possibly indicates that deviation from this sequence is poorly tolerated. The few SNPs observed are predicted to be pathogenic and all increase or decrease the number of UG in the sequence (VarSome). Further work will be needed to define whether any of these SNPs can be exploited to inform on disease onset or progression. Taken together, our data reinforce the evolutionary importance of TDP-43-mediated regulation of G3BP1, and by extension, its cellular function.

Transcriptomics data indicate that G3BP1 is expressed at relatively low levels in neurons (GTEx) and this could suggest that it is subject to very fine/tight regulation. We show that the short 3’UTR *G3BP1* transcript is the predominant one expressed in disease-vulnerable frontal cortex and lumbar spinal motor neurons and *G3BP1* mRNA levels were reduced in ALS/FTLD neurons bearing TDP-43 cytoplasmic inclusions/nuclear clearance. Our data in axotomized motor neurons, where loss of nuclear TDP-43 positively correlated with a sharp decrease in total G3BP1 levels is supportive of this possibility. Finally, near-physiological expression of ALS-associated TDP-43 mutations had the same impact on G3BP1 levels as nuclear TDP-43 depletion suggesting that TDP-43 mutations can act through a loss of function mechanism.

G3BP1 is essential for neuronal homeostasis and survival (Zekri *et al*., 2005; Martin *et al*., 2013; Martin *et al*., 2016b; Sahoo *et al*., 2018). Indeed, *G3BP1* genetic deletion in *129/sv* mice results in widespread neuronal loss, while the same deletion on a *Balb/c* background yields synaptic and locomotor deficits, with 75% of homozygotes displaying paralysis of at least one limb (Zekri *et al*., 2005; Martin *et al*., 2013). In addition, these mice have impairments in neuronal plasticity and in calcium/glutamate signaling (Martin *et al*., 2013; Martin *et al*., 2016a). G3BP1 is also a known target of certain enteroviruses, such as poliovirus and coxsackievirus B3, which each encode a protease that cleaves G3BP1 and effectively cripples the stress granule response in order to facilitate virus replication in motor neurons (White *et al*., 2007; Fung *et al*., 2013; Lloyd, 2016). It is noteworthy that poliovirus exhibits tropism for ALS-vulnerable regions (*i.e*. it infects the anterior horns of the spinal cord, brainstem and the motor cortex but not the oculomotor nerve) (Nagata *et al*., 2004; White *et al*., 2007). As lytic viruses, they cause selective motor neuron death and, in the case of poliovirus, induce a progressive paralysis (post-poliomyelitis syndrome) similar to that experienced by ALS patients (Xue *et al*., 2018).

Collectively, our data support that under normal conditions, TDP-43 functions to stabilize *G3BP1* transcripts via a highly conserved *cis* 3’UTR regulatory element. In addition, we observed reduced *G3BP1* mRNA levels in neurons bearing the TDP-43 pathological signature of ALS/FTLD. The loss of *G3BP1* mRNA, and thus the encoded protein that is essential to stress granule assembly, would preclude the launch of the protective neuronal response known as stress granule formation and draws into question the model that TDP-43 inclusions derive from defective stress granule disassembly. As we have previously published that deficient stress granule assembly contributes to enhanced neuronal vulnerability (McDonald *et al*., 2011; Aulas *et al*., 2012; Aulas *et al*., 2015), our data are consistent with the idea that stabilization of *G3BP1* mRNA and maintenance of stress granule dynamics could be a valid therapeutic avenue.

## ACKNOWLEDGEMENTS

We thank Nathalie Arbour for access to the plate reader and Aurélie Cleret-Buhot of the CRCHUM Cytometry, Cell Imaging and Molecular Pathology platform for help with microscopy. We thank Don Cleveland, Tania Kastelic and Dominique Cheneval for important discussions; Sarah Peyrard for lab management, and Julie Veriepe and Audrey Labarre for advice with *C. elegans* experiments. We thank André Dagenais, Yves Berthiaume and Francis Migneault for providing the pTRE-Tight vector and Marc Bilodeau for HepG2 cells. We thank Marjorie Labrecque and Jay Ross for help with RNA-seq analysis. We thank CVV present and past lab members for constructive and helpful feedback.

## FUNDING

This work was supported by the ALS Canada/Brain Canada Hudson Translational Team Grant (CVV), CIHR (JAP, JR and PL), the Muscular Dystrophy Association (CVV), the James Hunter ALS Initiative (JR, LZ), the ALS Association Milton Safenowitz Postdoctoral Fellowship (PMM), and the National Institutes of Health (National Institute for Neurological Disorders and Stroke R01-NS097542 (SJB), National Institute for Aging P30 AG053760 (SJB)), the University of Michigan Protein Folding Disease Initiative, and Ann Arbor Active Against ALS. HS is supported by an FRQS Doctoral Studentship. MT, CVV and JAP are FRQS Research Scholars.

## AUTHOR CONTRIBUTIONS

HS was involved in the project design, performed and analyzed most experiments, assembled the data, and wrote the manuscript with CVV. YK performed axotomies. GDT and PL performed and analyzed the EMSA. LD helped with luciferase assays. LD and AA performed the RNA-IPs. LZ consented patients. SX, PMM and JR performed and analyzed experiments in human tissues. JED helped with TDP-43^ΔNLS^ immunoblotting. EB, SJB and NBG provided iPSC-derived neurons and analyzed RNA-seq for polyA site usage. MT analyzed RNA-seq. JAP provided worms. ZM provided SH-SY5Y TDP-43^N352S/N352S^ cells. CVV designed and directed the project. All authors reviewed and approved the manuscript prior to submission.

## COMPETING INTERESTS

The authors have no competing interests to declare.

## SUPPLEMENTARY MATERIAL

Supplemental Figure.1: Expression of TDP-43 contructs.

Supplemental Figure 2: TDP-43 depletion does not influence *G3BP1* promotor activity.

Supplemental Figure 3: FUS expression in Fluorogold-positive injured motor neurons.

Supplementary Table I: Evolutionary conservation of the identified TDP-43 binding site in *G3BP1*.

Supplementary Table II. Demographic and clinical data from sporadic ALS and ALS/FTLD cases screened for G3BP1 mRNA puncta.

## Notes

### Competing Interest Statement

The authors have declared no competing interest.

## REFERENCES

Afroz T, Hock E-M, Ernst P, Foglieni C, Jambeau M, Gilhespy LAB, et al. Functional and dynamic polymerization of the ALS-linked protein TDP-43 antagonizes its pathologic aggregation. Nature Communications 2017; 8(1): 45.

Al-Chalabi A, Hardiman O. The epidemiology of ALS: a conspiracy of genes, environment and time. Nat Rev Neurol 2013; 9(11): 617–28.

Anders S, Pyl PT, Huber W. HTSeq--a Python framework to work with high-throughput sequencing data. Bioinformatics 2015; 31(2): 166–9.

Anderson P, Kedersha N. RNA granules. JCell Biol 2006; 172(6): 803–8.

Anderson P, Kedersha N. Stress granules: the Tao of RNA triage. Trends BiochemSci 2008; 33(3): 141–50.

Aulas A, Caron G, Gkogkas CG, Mohamed NV, Destroismaisons L, Sonenberg N, et al. G3BP1 promotes stress-induced RNA granule interactions to preserve polyadenylated mRNA. The Journal of cell biology 2015; 209(1): 73–84.

Aulas A, Stabile S, Vande Velde C. Endogenous TDP-43, but not FUS, contributes to stress granule assembly via G3BP. MolNeurodegener 2012; 7: 54.

Bailey TL, Boden M, Buske FA, Frith M, Grant CE, Clementi L, et al. MEME Suite: tools for motif discovery and searching. Nucleic Acids Research 2009; 37(suppl_2): W202–W8.

Baird T, Hogg J. Using tet-off cells and RNAi knockdown to assay mRNA decay. 2018. p. 161–73.

Boeynaems S, Bogaert E, Kovacs D, Konijnenberg A, Timmerman E, Volkov A, et al. Phase Separation of C9orf72 Dipeptide Repeats Perturbs Stress Granule Dynamics. Mol Cell 2017; 65(6): 1044–55.e5.

Buratti E, Baralle FE. Characterization and functional implications of the RNA binding properties of nuclear factor TDP-43, a novel splicing regulator of CFTR exon 9. JBiolChem 2001; 276(39): 36337–43.

Busskamp V, Lewis NE, Guye P, Ng AH, Shipman SL, Byrne SM, et al. Rapid neurogenesis through transcriptional activation in human stem cells. Molecular systems biology 2014; 10: 760.

Cerbini T, Funahashi R, Luo Y, Liu C, Park K, Rao M, et al. Transcription activator-like effector nuclease (TALEN)-mediated CLYBL targeting enables enhanced transgene expression and one-step generation of dual reporter human induced pluripotent stem cell (iPSC) and neural stem cell (NSC) lines. PloS one 2015; 10(1): e0116032.

Colombrita C, Onesto E, Megiorni F, Pizzuti A, Baralle FE, Buratti E, et al. TDP-43 and FUS RNA-binding proteins bind distinct sets of cytoplasmic messenger RNAs and differently regulate their post-transcriptional fate in motoneuron-like cells. The Journal of biological chemistry 2012; 287(19): 15635–47.

Costessi L, Porro F, Iaconcig A, Muro AF. TDP-43 regulates beta-adducin (Add2) transcript stability. RNA biology 2014; 11(10): 1280–90.

Deshaies JE, Shkreta L, Moszczynski AJ, Sidibe H, Semmler S, Fouillen A, et al. TDP-43 regulates the alternative splicing of hnRNP A1 to yield an aggregation-prone variant in amyotrophic lateral sclerosis. Brain 2018; 141(5): 1320–33.

Dewey CM, Cenik B, Sephton CF, Dries DR, Mayer P, 3rd, Good SK, et al. TDP-43 is directed to stress granules by sorbitol, a novel physiological osmotic and oxidative stressor. Mol Cell Biol 2011; 31(5): 1098–108.

Dewey CM, Cenik B, Sephton CF, Dries DR, Mayer P, III, Good SK, et al. TDP-43 is directed to stress granules by sorbitol, a novel physiological osmotic and oxidative stressor. MolCell Biol 2010; 31(5): 1098–108.

Di Tomasso G, Lampron P, Dagenais P, Omichinski JG, Legault P. The ARiBo tag: a reliable tool for affinity purification of RNAs under native conditions. Nucleic acids research 2011; 39(3): e18–e.

Fernandopulle MS, Prestil R, Grunseich C, Wang C, Gan L, Ward ME. Transcription Factor-Mediated Differentiation of Human iPSCs into Neurons. Curr Protoc Cell Biol 2018; 79(1): e51.

Fiesel FC, Voigt A, Weber SS, Van den HC, Waldenmaier A, Gorner K, et al. Knockdown of transactive response DNA-binding protein (TDP-43) downregulates histone deacetylase 6. EMBO J 2010; 29(1): 209–21.

Forman MS, Trojanowski JQ, Lee VMY. TDP-43: a novel neurodegenerative proteinopathy. Curr Opin Neurobiol 2007; 17(5): 548–55.

Fung G, Ng CS, Zhang J, Shi J, Wong J, Piesik P, et al. Production of a Dominant-Negative Fragment Due to G3BP1 Cleavage Contributes to the Disruption of Mitochondria-Associated Protective Stress Granules during CVB3 Infection. PloS one 2013; 8(11): e79546.

Gu J, Wu F, Xu W, Shi J, Hu W, Jin N, et al. TDP-43 suppresses tau expression via promoting its mRNA instability. Nucleic Acids Res 2017; 45(10): 6177–93.

Guillén-Boixet J, Kopach A, Holehouse AS, Wittmann S, Jahnel M, Schlüßler R. et al. RNA-Induced Conformational Switching and Clustering of G3BP Drive Stress Granule Assembly by Condensation. Cell 2020; 181(2): 346–61.e17.

Ha KCH, Blencowe BJ, Morris Q. QAPA: a new method for the systematic analysis of alternative polyadenylation from RNA-seq data. Genome Biol 2018; 19(1): 45.

Hui J, Hung LH, Heiner M, Schreiner S, Neumüller N, Reither G, et al. Intronic CA repeat and CA rich elements: a new class of regulators of mammalian alternative splicing. The EMBO journal 2005; 24(11): 1988–98.

Humoud MN, Doyle N, Royall E, Willcocks MM, Sorgeloos F, van Kuppeveld F, et al. Feline Calicivirus Infection Disrupts Assembly of Cytoplasmic Stress Granules and Induces G3BP1 Cleavage. J Virol 2016; 90(14): 6489–501.

Kabashi E, Lin L, Tradewell ML, Dion PA, Bercier V, Bourgouin P, et al. Gain and loss of function of ALS-related mutations of TARDBP (TDP-43) cause motor deficits in vivo. HumMolGenet 2010; 19(4): 671–83.

Khalfallah Y, Kuta R, Grasmuck C, Prat A, Durham HD, Vande Velde C. TDP-43 regulation of stress granule dynamics in neurodegenerative disease-relevant cell types. Sci Rep 2018; 8(1): 7551.

Kim D, Paggi JM, Park C, Bennett C, Salzberg SL. Graph-based genome alignment and genotyping with HISAT2 and HISAT-genotype. Nat Biotechnol 2019; 37(8): 907–15.

Krach F, Batra R, Wheeler EC, Vu AQ, Wang R, Hutt K, et al. Transcriptome-pathology correlation identifies interplay between TDP-43 and the expression of its kinase CK1E in sporadic ALS. Acta neuropathologica 2018; 136(3): 405–23.

Kuo PH, Chiang CH, Wang YT, Doudeva LG, Yuan HS. The crystal structure of TDP-43 RRM1-DNA complex reveals the specific recognition for UG- and TG-rich nucleic acids. Nucleic Acids Res 2014; 42(7): 4712–22.

Lam RS, Topfer FM, Wood PG, Busskamp V, Bamberg E. Functional Maturation of Human Stem Cell-Derived Neurons in Long-Term Cultures. PloS one 2017a; 12(1): e0169506.

Lam RS, Töpfer FM, Wood PG, Busskamp V, Bamberg E. Functional Maturation of Human Stem Cell-Derived Neurons in Long-Term Cultures. PloS one 2017b; 12(1): e0169506.

Lau AG, Irier HA, Gu J, Tian D, Ku L, Liu G, et al. Distinct 3’UTRs differentially regulate activity-dependent translation of brain-derived neurotrophic factor (BDNF). Proc Natl Acad Sci U S A 2010; 107(36): 15945–50.

Liu-Yesucevitz L, Bilgutay A, Zhang YJ, Vanderwyde T, Citro A, Mehta T, et al. Tar DNA binding protein-43 (TDP-43) associates with stress granules: analysis of cultured cells and pathological brain tissue. PLoSOne 2010; 5(10): e13250.

Lloyd RE. Enterovirus Control of Translation and RNA Granule Stress Responses. Viruses 2016; 8(4): 93–.

Love MI, Huber W, Anders S. Moderated estimation of fold change and dispersion for RNA-seq data with DESeq2. Genome Biol 2014; 15(12): 550.

Lukavsky PJ, Daujotyte D, Tollervey JR, Ule J, Stuani C, Buratti E, et al. Molecular basis of UG-rich RNA recognition by the human splicing factor TDP-43. Nat Struct Mol Biol 2013; 20(12): 1443–9.

Mackenzie IR, Nicholson AM, Sarkar M, Messing J, Purice MD, Pottier C, et al. TIA1 Mutations in Amyotrophic Lateral Sclerosis and Frontotemporal Dementia Promote Phase Separation and Alter Stress Granule Dynamics. Neuron 2017; 95(4): 808–16 e9.

Martin S, Bellora N, Gonzalez-Vallinas J, Irimia M, Chebli K, de Toledo M, et al. Preferential binding of a stable G3BP ribonucleoprotein complex to intron-retaining transcripts in mouse brain and modulation of their expression in the cerebellum. J Neurochem 2016a; 139(3): 349–68.

Martin S, Bellora N, González-Vallinas J, Irimia M, Chebli K, Toledo M, et al. Preferential binding of a stable G3BP ribonucleoprotein complex to intron-retaining transcripts in mouse brain and modulation of their expression in the cerebellum. Journal of Neurochemistry 2016b; 139(3): 349–68.

Martin S, Zekri L, Metz A, Maurice T, Chebli K, Vignes M, et al. Deficiency of G3BP1, the stress granules assembly factor, results in abnormal synaptic plasticity and calcium homeostasis in neurons. J Neurochem 2013; 125(2): 175–84.

McDonald KK, Aulas A, Destroismaisons L, Pickles S, Beleac E, Camu W, et al. TAR DNA-binding protein 43 (TDP-43) regulates stress granule dynamics via differential regulation of G3BP and TIA-1. HumMolGenet 2011; 20(7): 1400–10.

Melamed Z, Lopez-Erauskin J, Baughn MW, Zhang O, Drenner K, Sun Y, et al. Premature polyadenylation-mediated loss of stathmin-2 is a hallmark of TDP-43-dependent neurodegeneration. Nature neuroscience 2019; 22(2): 180–90.

Moisse K, Volkening K, Leystra-Lantz C, Welch I, Hill T, Strong MJ. Divergent patterns of cytosolic TDP-43 and neuronal progranulin expression following axotomy: implications for TDP-43 in the physiological response to neuronal injury. Brain Res 2009; 1249: 202–11.

Mompean M, Romano V, Pantoja-Uceda D, Stuani C, Baralle FE, Buratti E, et al. The TDP-43 N-terminal domain structure at high resolution. Febs j 2016; 283(7): 1242–60.

Nagata N, Iwasaki T, Ami Y, Sato Y, Hatano I, Harashima A, et al. A poliomyelitis model through mucosal infection in transgenic mice bearing human poliovirus receptor, TgPVR21. Virology 2004; 321(1): 87–100.

Neumann M, Sampathu DM, Kwong LK, Truax AC, Micsenyi MC, Chou TT, et al. Ubiquitinated TDP-43 in frontotemporal lobar degeneration and amyotrophic lateral sclerosis. Science 2006; 314(5796): 130–3.

Paz I, Kosti I, Ares M, Jr., Cline M, Mandel-Gutfreund Y. RBPmap: a web server for mapping binding sites of RNA-binding proteins. Nucleic Acids Res 2014; 42(Web Server issue): W361–7.

Prudencio M, Belzil VV, Batra R, Ross CA, Gendron TF, Pregent LJ, et al. Distinct brain transcriptome profiles in C9orf72-associated and sporadic ALS. Nat Neurosci 2015; 18(8): 1175–82.

Rodriguez A, Perez-Gracia E, Espinosa JC, Pumarola M, Torres JM, Ferrer I. Increased expression of water channel aquaporin 1 and aquaporin 4 in Creutzfeldt-Jakob disease and in bovine spongiform encephalopathy-infected bovine-PrP transgenic mice. Acta neuropathologica 2006; 112(5): 573–85.

Romo L, Ashar-Patel A, Pfister E, Aronin N. Alterations in mRNA 3’ UTR Isoform Abundance Accompany Gene Expression Changes in Human Huntington’s Disease Brains. Cell reports 2017; 20(13): 3057–70.

Sahoo PK, Lee SJ, Jaiswal PB, Alber S, Kar AN, Miller-Randolph S, et al. Axonal G3BP1 stress granule protein limits axonal mRNA translation and nerve regeneration. Nature communications 2018; 9(1): 3358–.

Salvail-Lacoste A, Di Tomasso G, Piette BL, Legault P. Affinity purification of T7 RNA transcripts with homogeneous ends using ARiBo and CRISPR tags. RNA (New York, NY) 2013; 19(7): 1003–14.

Sanders DW, Kedersha N, Lee DSW, Strom AR, Drake V, Riback JA, et al. Competing Protein-RNA Interaction Networks Control Multiphase Intracellular Organization. Cell 2020; 181(2): 306–24.e28.

Sidibé H, Vande Velde C. RNA Granules and Their Role in Neurodegenerative Diseases. In: Oeffinger M, Zenklusen D, editors. The Biology of mRNA: Structure and Function. Cham: Springer International Publishing; 2019. p. 195–245.

Tank EM, Figueroa-Romero C, Hinder LM, Bedi K, Archbold HC, Li X, et al. Abnormal RNA stability in amyotrophic lateral sclerosis. Nature Communications 2018; 9(1): 2845.

Taylor JP, Brown RH, Jr., Cleveland DW. Decoding ALS: from genes to mechanism. Nature 2016; 539(7628): 197–206.

Tourriere H, Chebli K, Zekri L, Courselaud B, Blanchard JM, Bertrand E, et al. The RasGAP-associated endoribonuclease G3BP assembles stress granules. JCell Biol 2003; 160(6): 823–31.

Tushev G, Glock C, Heumuller M, Biever A, Jovanovic M, Schuman EM. Alternative 3’ UTRs Modify the Localization, Regulatory Potential, Stability, and Plasticity of mRNAs in Neuronal Compartments. Neuron 2018; 98(3): 495–511.e6.

Urushitani M, Ezzi SA, Julien JP. Therapeutic effects of immunization with mutant superoxide dismutase in mice models of amyotrophic lateral sclerosis. ProcNatlAcadSciUSA 2007; 104(7): 2495–500.

Vaccaro A, Tauffenberger A, Ash PE, Carlomagno Y, Petrucelli L, Parker JA. TDP-1/TDP-43 regulates stress signaling and age-dependent proteotoxicity in Caenorhabditis elegans. PLoSGenet 2012; 8(7): e1002806.

Van Nostrand EL, Freese P, Pratt GA, Wang X, Wei X, Xiao R, et al. A large-scale binding and functional map of human RNA-binding proteins. Nature 2020; 583(7818): 711–9.

Wang IF, Wu LS, Chang HY, Shen CK. TDP-43, the signature protein of FTLD-U, is a neuronal activity-responsive factor. JNeurochem 2008; 105(3): 797–806.

Weskamp K, Tank EM, Miguez R, Gómez NB, White M, Lin Z, et al. Neuronal hyperexcitability drives TDP43 pathology by upregulating shortened TDP43 protein isoforms. 2019: 648477.

White JP, Cardenas AM, Marissen WE, Lloyd RE. Inhibition of cytoplasmic mRNA stress granule formation by a viral proteinase. Cell HostMicrobe 2007; 2(5): 295–305.

Winton MJ, Igaz LM, Wong MM, Kwong LK, Trojanowski JQ, Lee VM. Disturbance of nuclear and cytoplasmic TAR DNA-binding protein (TDP-43) induces disease-like redistribution, sequestration, and aggregate formation. The Journal of biological chemistry 2008; 283(19): 13302–9.

Xiao S, Sanelli T, Chiang H, Sun Y, Chakrabartty A, Keith J, et al. Low molecular weight species of TDP-43 generated by abnormal splicing form inclusions in amyotrophic lateral sclerosis and result in motor neuron death. Acta neuropathologica 2015; 130(1): 49–61.

Xue YC, Feuer R, Cashman N, Luo H. Enteroviral Infection: The Forgotten Link to Amyotrophic Lateral Sclerosis? Frontiers in molecular neuroscience 2018; 11: 63–.

Yang P, Mathieu C, Kolaitis R-M, Zhang P, Messing J, Yurtsever U, et al. G3BP1 Is a Tunable Switch that Triggers Phase Separation to Assemble Stress Granules. Cell 2020; 181(2): 325–45.e28.

Yu J, Chau KF, Vodyanik MA, Jiang J, Jiang Y. Efficient feeder-free episomal reprogramming with small molecules. PloS one; 2011. p. e17557.

Zekri L, Chebli K, Tourriere H, Nielsen FC, Hansen TV, Rami A, et al. Control of fetal growth and neonatal survival by the RasGAP-associated endoribonuclease G3BP. MolCell Biol 2005; 25(19): 8703–16.

Zhang Y, Pak C, Han Y, Ahlenius H, Zhang Z, Chanda S, et al. Rapid single-step induction of functional neurons from human pluripotent stem cells. Neuron 2013; 78(5): 785–98.

Zheng D, Wang R, Ding Q, Wang T, Xie B, Wei L, et al. Cellular stress alters 3’UTR landscape through alternative polyadenylation and isoform-specific degradation. Nature Communications 2018; 9(1): 2268.

