## Supplemental material for "TDP-43 stabilizes transcripts encoding stress granule protein G3BP1: potential relevance to ALS/FTD"

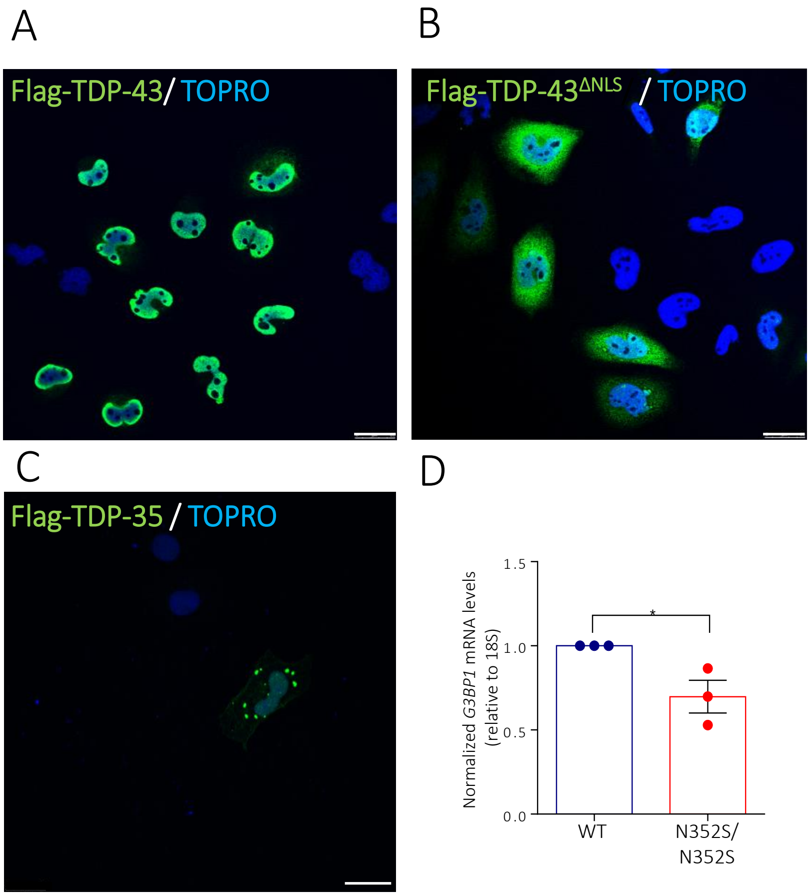


**Supplementary figure 1. Expression of TDP-43 constructs.** (A) HeLa cells were treated with siTDP-43 or siControl, then transfected with Flag-TDP-43 (WT) or (B) Flag-TDP-43$^{\Delta NLS}$ or (C) only transfected with Flag-TDP-35 and immunolabelled with anti-Flag and TOPRO. (D) mRNA from SH-SY5Y WT and TDP-43 N352S/N352S cells were extracted, and RT-qPCR was performed to evaluate *G3BP1* transcript levels. Data from 3 independent experiments are expressed as the mean fold change ± SEM; Unpaired t test $^{*}p<0.05$ Scale bar, 25µm.

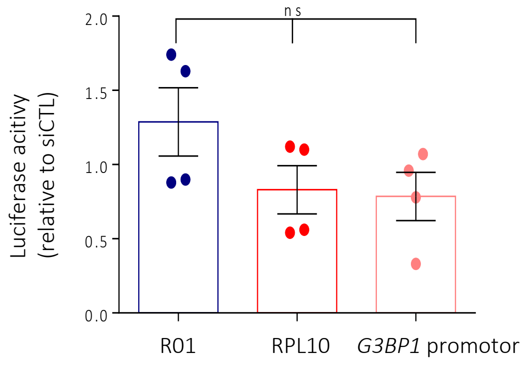


**Supplementary figure 2. TDP-43 depletion does not influence *G3BP1* promotor activity.** HeLa cells were transfected with siTDP-43 or siControl, then co-transfected with the indicated reporter plasmids. Luciferase activity of G3BP1 promotor is expressed relative to siControl cells. RPL10 and R01 (random sequences) are used as controls, mean ± SEM, n=4. Data from 4 independent experiments are expressed as the mean fold change ± SEM; Unpaired t test

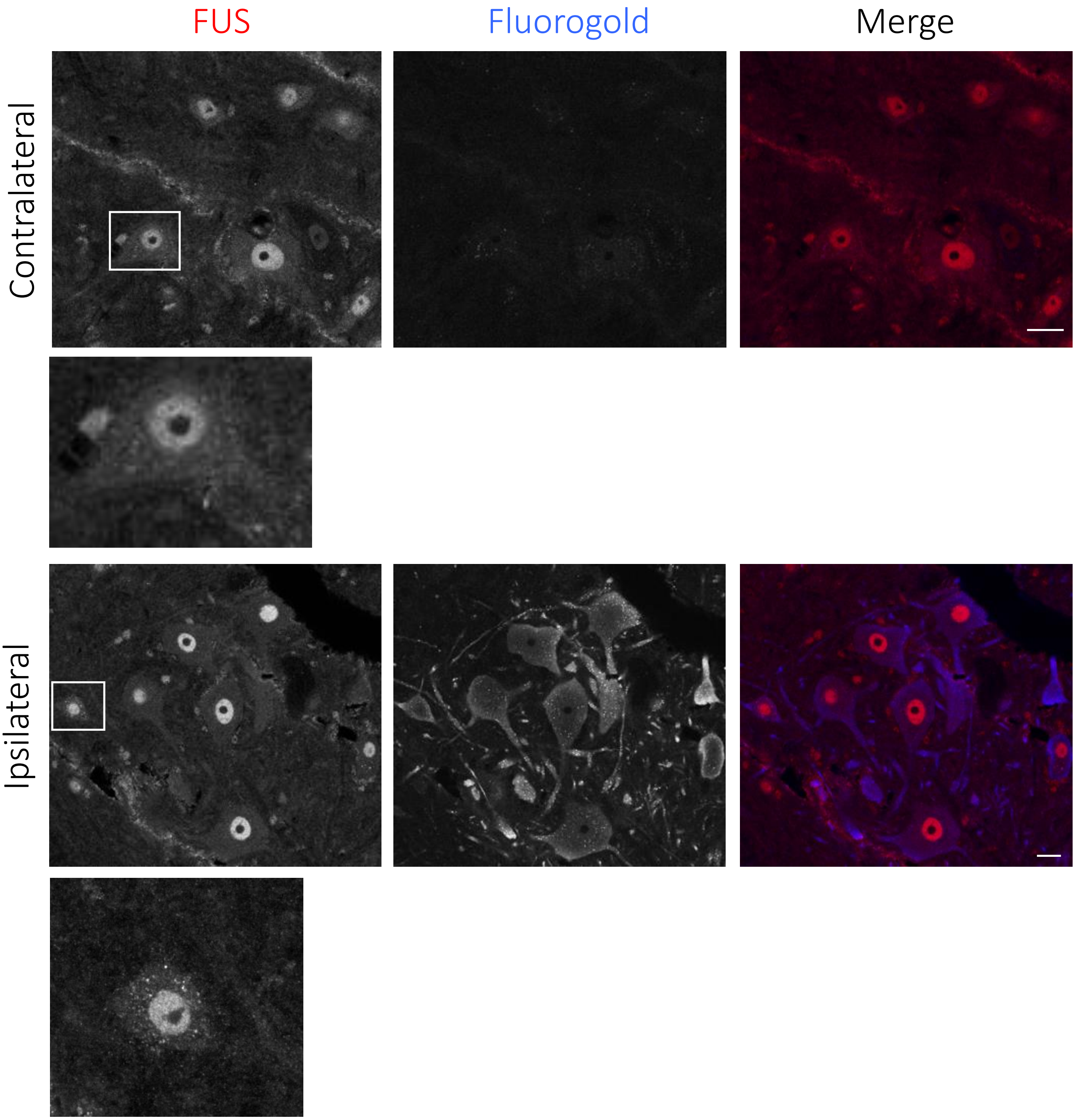


**Supplementary figure 3. FUS expression in Fluorogold-positive injured motor neurons.** Representative images of FUS expression (red) in Fluorogold-positive (blue) neurons on the ipsilateral side and Fluorogold-negative in the contralateral of the ventral spinal cord of axotomized mice at day 7. Scale bar, 10 µm.

Supplementary Table I. Evolutionary conservation of the identified TDP-43 binding site in *G3BP1*. Conservation of the regulatory element in 13 species and the homology between their respective 3'UTR sequences and the conserved elements.

| Species | Sequence | 3'UTR homology to human (%) | Local homology to human (%) |
|---|---|---|---|
| *Homo sapiens* | UGUGUUAAUGGUGUGU | NA | NA |
| *Pongo abelli* (orangutan) | UGUGUUAAUGGUGUGU | 98% | 100% |
| *Macaca mulatta* (Macaque) | UGUGUUAAUGGUGUGU | 96% | 100% |
| *Ictidomys tridecemlineatus* (Thriteen-lined ground squirrel) | UGUGUUAAUGGUGUGU | 87% | 100% |
| *Sus scrafa* (Pig) | UGUGUUAAUGGUGUGU | 83% | 100% |
| *Rattus norvegicus* (Rat) | UGUGUUCAUGGUGUGU | 77% | 95% |
| *Mus musculus* (Mouse) | UGUGUUCAUGGUGUGU | 76% | 95% |
| *Mesocricetus auratus* (Golden Hamster) | UGUGUUCAUGGUGUGU | 61% | 95% |
| *Cricetulus griseus* (Chinese Hamster) | UGUGUUCAUGGUGUGU | 0% | 95% |
| *Falco peregrinus* (Falcon) | UUUGUUUGUGUUAUGG | 0% | 87% |
| *Anas platyrhynchos* (Mallard) | UUUGUUUGUGUUAUGG | 0% | 87% |
| *Xenopus laevis* (clawed frog) | UGUCGUGACAGACUGU | 0% | 75% |
| *Caenorhabditis elegans* | UAUUUAUUGUUU | 0% | 67% |

Supplementary Table II. Demographic and clinical data from sporadic ALS and ALS/FTLD cases screened for G3BP1 mRNA puncta.

| Case | Sex | Age (at autopsy) | Diagnosis | TDP43 positive* |
|---|---|---|---|---|
| 1 | M | 70 | sporadic ALS | No |
| 2 | F | 57 | sporadic ALS | No |
| 3 | M | 37 | sporadic ALS | No |
| 4 | M | 73 | sporadic ALS | No |
| 5 | M | 74 | ALS/FTLD | Yes |
| 6 | F | 66 | ALS/FTLD | Yes |
| 7 | F | 57 | ALS/FTLD | Yes |
| 8 | M | 73 | ALS/FTLD | Yes |

* case is positive for TDP-43 pathology in the orbitofrontal cortex (BA11/47)
